# Multi-omics characterization of extracellular vesicles derived from virus-positive Merkel cell carcinoma cells

**DOI:** 10.64898/2026.08.15.745013

**Authors:** Ute Andrea Westerkamp, Patrick Blümke, Amanda Salviano-Silva, Claudia Schmidt, Thomas Mair, Bente Siebels, Jiabin Huang, Nicole Fischer

## Abstract

Merkel cell carcinoma (MCC) is a highly aggressive skin cancer, with approximately 80% of cases driven by Merkel cell polyomavirus (MCPyV). Although extracellular vesicles (EVs) are increasingly recognized as mediators of intercellular communication within the tumor microenvironment, their molecular cargo in MCPyV-positive MCC has not been comprehensively characterized. Here, we performed a multi-omics characterization of EVs released by two MCPyV-positive MCC cell lines. EVs were isolated by differential ultracentrifugation and characterized by nanoparticle tracking analysis, imaging flow cytometry, cryo-electron microscopy, and immunoblotting, demonstrating a heterogeneous population of small and large EVs. Proteomic and transcriptomic analyses revealed that MCC-derived EVs possess distinct protein, mRNA, and miRNA cargo compared with their parental cells, with enrichment of molecules associated with gene expression, RNA processing, intracellular signaling, and vesicle-mediated transport. Despite differences in the molecular composition of EVs derived from WaGa and MKL-1 cells, functional enrichment analyses revealed highly similar biological pathways. To investigate whether the viral oncoprotein small T antigen (sT) contributes to EV cargo composition, EVs from inducible sT knockdown cells were analyzed. Loss of sT was associated with modest changes in the EV proteome and mRNA cargo, whereas the overall EV-associated miRNA profile remained largely unchanged. Collectively, these findings provide the first comprehensive molecular characterization of EVs released by MCPyV-positive MCC cells and establish a foundation for investigating the contribution of EV-mediated communication to MCC biology and tumor–microenvironment interactions.

## Introduction

Merkel cell carcinoma (MCC) is a rare but highly aggressive neuroendocrine skin cancer characterized by a high propensity for metastasis and recurrence (1). Although immune checkpoint inhibitors have improved clinical outcomes, therapeutic options remain limited, underscoring the need for a better understanding of the molecular mechanisms driving MCC tumorigenesis (2, 3). Approximately 80% of MCCs are associated with Merkel cell polyomavirus (MCPyV), whereas the remaining tumors are driven by ultraviolet (UV)-induced mutagenesis (4–7). A defining feature of MCPyV-positive MCC is the clonal integration of the viral genome into the host genome, resulting in constitutive expression of the viral oncoproteins small T antigen (sT) and a truncated large T antigen (LT) (5, 8). Whereas truncated LT is essential for tumor maintenance, sT is considered the principal oncogenic driver of virus-positive MCC and is sufficient to transform cells in experimental models (9–12). Through interactions with numerous cellular proteins, sT regulates pathways involved in proliferation, migration, metastasis, and immune evasion (13–18). Furthermore, sT remodels the host transcriptome and cell-surface proteome, including upregulation of the immune checkpoint molecule CD47, thereby promoting immune escape (19, 20). These observations suggest that sT broadly influences the molecular phenotype of MCC cells and raise the possibility that it also contributes to shaping the molecular cargo of extracellular vesicles (EVs).

Extracellular vesicles (EVs) are lipid bilayer-enclosed particles released by virtually all cell types that mediate intercellular communication through the transfer of proteins, lipids, and nucleic acids (21–23). Although EV cargo is influenced by the molecular composition and physiological state of the parental cell, numerous studies have shown that proteins and RNAs can become enriched in EVs relative to their parental cells, indicating that EV cargo does not simply reflect the cellular proteome or transcriptome (22–25). Consequently, EV cargo does not necessarily mirror the cellular proteome or transcriptome and can elicit specific responses in recipient cells. Within the tumor microenvironment, tumor-derived EVs contribute to immune evasion, extracellular matrix remodeling, angiogenesis, metastasis, and therapy resistance (26, 27). Recent work further demonstrated that MCPyV virions can associate with EVs released from infected primary dermal fibroblasts, thereby enhancing infection efficiency and reducing susceptibility to antibody-mediated neutralization. Proteomic analyses further indicated that MCPyV infection is associated with changes in EV protein composition, including proteins involved in antiviral and immune-regulatory processes (28). Because EV proteins and RNAs can act in concert to influence recipient cells, comprehensive multi-omics analyses are increasingly being used to define EV-mediated communication within the tumor microenvironment (27, 29–31). In MCC, EV-mediated transfer of miR-375 promotes fibroblast polarization toward a cancer-associated phenotype through repression of RBPJ and TP53, providing evidence that EV cargo can actively contribute to MCC tumorigenesis (32, 33). Despite growing evidence that EVs participate in both MCPyV infection and MCC progression, a comprehensive molecular characterization integrating the proteome, mRNA cargo, and miRNA cargo of EVs released by MCPyV-positive MCC cells is currently lacking. Furthermore, it remains unknown whether the viral oncoprotein small T antigen (sT) influences EV cargo composition. Because EV-mediated communication is driven by the coordinated action of proteins and coding and non-coding RNAs, integrated multi-omics analyses are required to comprehensively define EV cargo composition and its potential biological functions.

In this study, we performed an integrated multi-omics characterization of extracellular vesicles released by two independent MCPyV-positive MCC cell lines. Using quantitative proteomics, RNA sequencing, and small RNA sequencing, we compared the protein and RNA cargo of EVs with that of their corresponding parental cells and investigated how inducible depletion of the viral small T antigen affects EV cargo composition. Together, these analyses provide the first comprehensive multi-omics characterization of EVs released by MCPyV-positive MCC cells and establish a foundation for future studies investigating EV-mediated communication in MCC.

## Methods

### Cell culture

WaGa (9) and MKL-1 (34) MCPyV-positive Merkel cell carcinoma (MCC) cell lines were cultured in RPMI 1640 medium supplemented with 10% fetal bovine serum (FBS), 1% penicillin–streptomycin (P/S), and 1% L-glutamine at 37 °C in a humidified incubator with 5% CO₂. For inducible knockdown of the viral small T antigen (sT), WaGa and MKL-1 cells carrying a doxycycline-inducible shRNA construct targeting sT or a scrambled control shRNA (20) were treated with 1 µg/mL doxycycline or an equivalent volume of DMSO for 3 days before further analyses as described before (20).

### Extracellular vesicle (EV) isolation

For EV isolation, cells were seeded at a density of 5 × 10⁵ cells/mL in 25–35 mL RPMI 1640 medium supplemented with 5% exosome-depleted FBS (Gibco, Cat. No. A27208-03), 1% penicillin– streptomycin, and 1% L-glutamine. Cells were cultured for 3 days in the presence of 1 µg/mL doxycycline or DMSO.

Conditioned medium was collected and subjected to sequential differential centrifugation to remove cells, cell debris, and large extracellular particles. Briefly, conditioned medium was centrifuged at 400 × *g* for 5 min, and the resulting cell pellet was retained for preparation of whole-cell lysates to verify sT knockdown by Western blotting. The supernatant was subsequently centrifuged at 1,000 × *g* for 10 min, 3,220 × *g* for 10 min, and 10,000 × *g* for 30 min at 4 °C.

The resulting supernatant was transferred to polypropylene ultracentrifuge tubes and centrifuged at 100,000 × *g* for 2 h at 4 °C using an SW40 Ti swinging-bucket rotor (Beckman Coulter). The EV pellet was washed once with phosphate-buffered saline (PBS) and subjected to a second ultracentrifugation step at 100,000 × *g* for 2 h. The final EV pellet was resuspended in PBS supplemented with protease inhibitor (Roche) and stored at −80 °C until further analysis.

EV isolation and characterization were performed in accordance with the Minimal Information for Studies of Extracellular Vesicles 2023 (MISEV2023) guidelines (35).

### RNA and small-RNA Sequencing

Total RNA was isolated from 200 µL of EV suspension using the miRNeasy Micro Kit (Qiagen, Germany) according to the manufacturer’s instructions. RNA concentration was determined using the Qubit™ Fluorometer and the Qubit™ RNA High Sensitivity (HS) Assay Kit (Thermo Fisher Scientific), while RNA purity was assessed using a NanoDrop 2000 spectrophotometer (PeqLab). RNA integrity was evaluated using either an Agilent 2100 Bioanalyzer or an Agilent TapeStation system (Agilent Technologies, USA).

Libraries for total RNA sequencing were first subjected to ribosomal RNA (rRNA) depletion using the RiboCop rRNA Depletion Kit (Lexogen, Austria). Subsequently, the RNA was processed via the CORALL Total RNA-Seq V2 Kit (Lexogen, Austria) according to the manufacturer’s protocols. For small RNA sequencing libraries were generated using the Small RNA-Seq Library Prep Kit (Lexogen, Austria) according to the manufacturers protocol. Library quality and fragment size distribution were assessed using an Agilent Bioanalyzer.

Sequencing was performed on an Illumina NextSeq 500 platform (Illumina, USA) using 75-bp reads. Total RNA libraries were sequenced to a depth of approximately 20–30 million reads per sample, whereas small RNA libraries yielded approximately 1–10 million reads per sample.

### Western Blotting

Three days after treatment with 1 µg/mL doxycycline or an equivalent volume of DMSO, conditioned medium was collected for EV isolation and the corresponding cell pellets were harvested for whole-cell lysate preparation. Cell pellets were washed once with ice-cold phosphate-buffered saline (PBS) and lysed in 5× Passive Lysis Buffer (Promega, USA). Protein concentrations of cell lysates were determined using the Qubit™ Fluorometer and the Qubit™ Protein Assay Kit (Thermo Fisher Scientific) to ensure equal protein loading.

For Western blot analysis, 20–30 µg of protein from whole-cell lysates or EV preparations was mixed with 4× protein loading buffer, denatured at 95 °C for 10 min, and separated by 12% SDS–PAGE. Proteins were transferred onto polyvinylidene difluoride (PVDF) membranes, which were subsequently blocked with 5% non-fat dry milk in PBS containing 0.05% Tween-20 (PBST).

Membranes were incubated overnight at 4 °C with the following primary antibodies: anti-AGO2 (MABE523, clone 11A9, Merck Millipore; 1:500), anti-ALIX (ABC40, Merck Millipore; 1:1,000), anti-CD9 (#13174, clone D8O1A, Cell Signaling Technology; 1:500), anti-CD81 (sc-166029, clone B-11, Santa Cruz Biotechnology; 1:500), anti-Flotillin-1 (610820, clone 18/Flotillin-1, BD Biosciences; 1:1,000), and anti-GM130 (610822, clone 35/GM130, BD Biosciences; 1:500).

After washing with PBST, membranes were incubated for 1–2 h at room temperature with horseradish peroxidase (HRP)-conjugated secondary antibodies: anti-mouse IgG-HRP (#12-371, Merck Millipore; 1:10,000) or anti-rabbit IgG-HRP (#12-348, Merck Millipore; 1:3,000). Immunoreactive bands were detected using Western Lightning Plus-ECL substrate (PerkinElmer) and imaged with an ImageQuant™ 800 imaging system (Cytiva). Band intensities were quantified using Fiji (ImageJ).

### Nanoparticle Tracking Analysis (NTA)

Nanoparticle tracking analysis (NTA) was performed using a NanoSight LM14 instrument (Malvern Panalytical, UK) to determine the size distribution and particle concentration of EV preparations. Prior to analysis, EV samples were diluted 1:50 in sterile-filtered phosphate-buffered saline (PBS). For each sample, five 30-s videos were recorded with the camera level set to 15 and the screen gain set to 1.0. Recordings were analyzed using NanoSight NTA software version 3.0 (Malvern Panalytical) with a detection threshold of 6 and a screen gain of 10.

### Imaging flow cytometry

Imaging flow cytometry (IFCM) was performed to characterize the surface marker profile of EVs. EV preparations (3–5 µL) were stained in sterile-filtered phosphate-buffered saline (PBS) supplemented with 8% EV-depleted fetal bovine serum (FBS). Samples were incubated for 45 min at room temperature in the dark with fluorophore-conjugated antibodies against CD81 (BioLegend, #349503; 1:3), CD9 (BioLegend, #312105; 1:30), CD63 (BioLegend, #353011; 1:3), or CD47 (Miltenyi Biotec, #130-123-875). Following incubation, unbound antibodies were removed by ultrafiltration using 300-kDa Nanosep centrifugal filter units. EVs were washed with PBS containing 2% EV-depleted FBS and resuspended in 30 µL of cold wash buffer prior to acquisition.

Data were acquired on an Amnis® ImageStreamX® Mark II imaging flow cytometer (Cytek, USA) using a 60× objective at low flow rate with an acquisition time of 60 s per sample. Fluorescence was detected using the following channels: channel 1 (VioBlue; 435–480 nm), channel 2 (FITC; 480–560 nm), channel 3 (PE; 560–595 nm), and channel 7 (Pacific Blue; 435–505 nm). Data analysis was performed using IDEAS software (version 6.2, Cytek), and an identical gating strategy was applied to all samples.

To minimize coincident particle detection (swarm detection), only events containing a single fluorescent spot were included in the analysis, while events containing multiple fluorescent spots were excluded.

### Cryogenic Electron-Microscopy

Cryogenic Electron-Microscopy (Cryo-EM) was performed to examine the morphology of EVs. EV preparations (approximately 10^9^ EVs/ml) were mixed with colloidal gold beads for 5-10 min. Holey carbon-coated copper grids were glow discharged immediately before sample application. Subsequently, 3µl of the EV suspension was applied to the grids allowed to adsorb for 30 sec, blotted with filter paper to remoce excess liquid, and vitrified by plunge freezing in an ethane-propane mixture using a Vitrobot Mark IV (Thermo Fisher Scientific, USA). Vitrified grids were maintained under liquid nitrogen and imaged at −175 °C using a Talos Arctica 200 kV transmission electron microscope equipped with a Falcon IVi direct electron detector (Thermo Fisher Scientific, USA). Images were acquired using the manufacturer’s acquisition software and subsequently analyzed with FEI image analysis software.

### Liquid Chromatography Tandem mass spectrometry (LC-MS/MS)-based Proteomics

Protein concentration of EV preparations was determined using the Pierce™ BCA Protein Assay Kit (Thermo Fisher Scientific, USA). For each sample, 20 µg of protein was adjusted to a final volume of 50 µL and processed using the single-pot solid-phase-enhanced sample preparation (SP3) protocol as previously described (36). Briefly, proteins were reduced with 10 mM dithiothreitol, alkylated with 20 mM iodoacetamide, and bound to carboxylate-modified paramagnetic beads (Sera-Mag SpeedBeads, GE Healthcare) in 70% acetonitrile using a 1:1 mixture of hydrophilic and hydrophobic beads. After incubation, beads were washed twice with 100% acetonitrile and twice with 70% ethanol. Proteins were digested overnight at 37 °C with sequencing-grade modified trypsin (Promega) in 5 mM ammonium bicarbonate. Tryptic peptides were subsequently rebound to the beads in 95% acetonitrile, washed twice with 100% acetonitrile, eluted with 2% dimethyl sulfoxide (DMSO) in 1% formic acid, and dried in a vacuum concentrator.

Peptides were separated using an Ultimate 3000 nano-UHPLC system (Thermo Fisher Scientific, USA) equipped with a C18 trap column (100 µm × 20 mm, 5 µm particle size, 100 Å pore size; Thermo Fisher Scientific) and an analytical C18 column (75 µm × 250 mm, 1.7 µm particle size, 130 Å pore size; Waters). Peptide separation was performed over an 80-min gradient using 0.1% formic acid in water (buffer A) and 0.1% formic acid in acetonitrile (buffer B), with the acetonitrile concentration increasing from 2% to 30% over 60 min.

Mass spectrometric analysis was performed on a Q Exactive Orbitrap or Fusion mass spectrometer (Thermo Fisher Scientific, USA) coupled to the nano-UHPLC system via nano-electrospray ionization (1800V). On the QExactive data were acquired in data-dependent acquisition mode using a Top15 method. Full MS scans were recorded over an *m/z* range of 400–1,200 at a resolution of 70,000 (*m/z* 200) with an automatic gain control (AGC) target of 1 × 10^6^ ions and a maximum injection time of 240 ms. The 15 most abundant precursor ions with charge states of +2 to +5 were isolated (2 *m/z* window) and fragmented by higher-energy collisional dissociation using a normalized collision energy of 25%. MS/MS spectra were acquired at a resolution of 17,500 with an AGC target of 1 × 10⁵ ions, a maximum injection time of 50 ms, and a dynamic exclusion time of 20 s. On the Fusion for each MS1 scan, ions were accumulated for a maximum of 120 milliseconds or until a charge density of 2 x 10^5^ ions (AGC Target) was reached. Fourier-transformation based mass analysis of the data from the orbitrap mass analyzer was performed covering a mass range of m/z 400 – 1,300 with a resolution of 120,000 at m/z = 200. Peptides with charge states between 2+ - 5+ above an intensity threshold of 1,000 were isolated within a m/z 1.6 isolation window in Top Speed mode for 3 seconds from each precursor scan and fragmented with a normalized collision energy of 30% using higher energy collisional dissociation (HCD). MS2 scanning was performed, using an ion trap mass analyzer at a rapid scan rate, covering a mass range starting at m/z 120 and accumulated for 60 ms or to an AGC target of 1 x 10^5^. Already fragmented peptides were excluded for 30 s.

### Proteomics data analysis

LC–MS/MS data were analyzed using the Sequest HT search engine implemented in Proteome Discoverer (V3.1.0638, Thermo Fisher Scientific) against the reviewed Homo sapiens database (v2023-11-08). Carbamidomethylation of cysteine residues was specified as a fixed modification, whereas oxidation of methionine and protein N-terminal acetylation were included as variable modifications. A maximum of two missed tryptic cleavages was allowed, and peptides comprising 6– 144 amino acids were considered for identification.

Peptide-spectrum matches and protein identifications were filtered to a false discovery rate (FDR) of ≤1% using a target-decoy strategy. Label-free protein quantification was performed using the Minora Feature Detector within Proteome Discoverer. Protein abundances were log₂-transformed and normalized by median-centering across samples before downstream statistical analyses. Data derived from different mass spectrometers were integrated using the BERT algorithm for batch effect reduction with standard settings (37). Statistical analysis was performed in Perseus (V 2.0.10.0). Student’s t-test were performed with Benjamini Hochberg FDR-correction.

### RNA Sequencing data analysis

A total of 40 RNA-seq libraries were processed using the nf-core/rnaseq pipeline (v1.3dev; DOI: 10.5281/zenodo.1400710) within the nf-core framework (38). Workflow execution was performed with Nextflow (v21.04.0) (39). Sequencing reads were aligned using STAR (v2.6.1d) (40) against a combined reference comprising the human genome (GRCh38) and the corresponding Merkel cell polyomavirus (MCPyV) genome (KJ128379.1 for WaGa samples and FJ173815.1 for MKL-1 samples). Gene-level read counts were generated using featureCounts (v2.0.1) (41). Low-count genes were filtered by retaining only those with a read count > 3 in at least three samples. Raw count data were normalized and variance-stabilized using the variance-stabilizing transformation (VST) from DESeq2 (v1.34.0) (42). Principal component analysis (PCA) was performed on VST-transformed data to assess sample clustering and reproducibility. Differential abundance analysis was carried out using DESeq2, and transcripts with an absolute log₂ fold change ≥ 1 and an adjusted *P* value ≤ 0.05 were considered significantly differentially abundant. The VST-transformed counts were used for downstream clustering and visualization.

### small RNA Sequencing data analysis

Raw FASTQ files generated by small RNA sequencing were processed using Cutadapt (v3.5) (43) to remove 3’adapter sequences introduced during library preparation with the LEXOGEN Small RNA-Seq Library Prep Kit (adapter sequence: TGGAATTCTCGGGTGCCAAGGAACTCCAGTCAC). Reads with an average Phred quality score below 20, reads shorter than 5 nucleotides after trimming, and reads containing ambiguous bases (N) were discarded.

High-quality trimmed reads were aligned and quantified using the exceRpt smallRNA-seq pipeline (version 4.6.3) (44). he pipeline utilizes a hierarchical mapping strategy against a bundled reference database (exceRptDB_v4), which integrates the human genome (GRCh38/hg38) and transcriptome annotations (GENCODE, miRBase, GtRNAdb, piRNABank, and circBase), alongside the NCBI UniVec database for contaminant filtering. Unmapped reads were subsequently aligned to exogenous miRNA/rRNA libraries and a comprehensive panel of exogenous genomes (bacteria, fungi, plants, protists, and viruses) to identify non-human sequences.

Raw miRNA count data were analyzed using DESeq2 (v1.34.0) (42). Low-count miRNAs were filtered by retaining only those with a total read count > 10 across all samples. Counts were normalized using the DESeq2 median-of-ratios method, and the regularized logarithm (rlog) transformation was applied for visualization. Principal component analysis (PCA), hierarchical clustering, and heatmaps were generated using rlog-transformed data. Differential abundance analysis was performed using DESeq2, and miRNAs with an absolute log₂ fold change ≥ 2 and an adjusted *P* value ≤ 0.05 were considered significantly differentially abundant. Volcano plots display DESeq2 differential abundance results.

Predicted biological functions of differentially abundant miRNAs were investigated by Gene Ontology enrichment analysis using experimentally validated target genes retrieved from miRTarBase. Target gene interaction networks were visualized using Cytoscape (45).

## Results

### WaGa Merkel cell carcinoma cells release a heterogeneous EV population displaying characteristics of both exosomes and microvesicles

To characterize extracellular vesicles (EVs) released by MCPyV-positive Merkel cell carcinoma (MCC) cells and to investigate the contribution of the viral small T antigen (sT) to EV cargo, we employed previously established WaGa and MKL-1 cell lines carrying a doxycycline-inducible shRNA system that enables conditional depletion of sT (20). EVs were isolated from conditioned medium of untreated cells as well as from control and sT knockdown cultures by differential centrifugation followed by ultracentrifugation (Figure 1A, Supplementary Figure S1). The isolated EVs were subsequently characterized with respect to their physical, biochemical, and morphological properties. Comparable results were obtained for EVs isolated from the independent MCPyV-positive MCC cell line MKL-1 (Figure 1B-D, Supplementary Figure S1), indicating that these characteristics are not restricted to WaGa cells.

**Figure 1:**
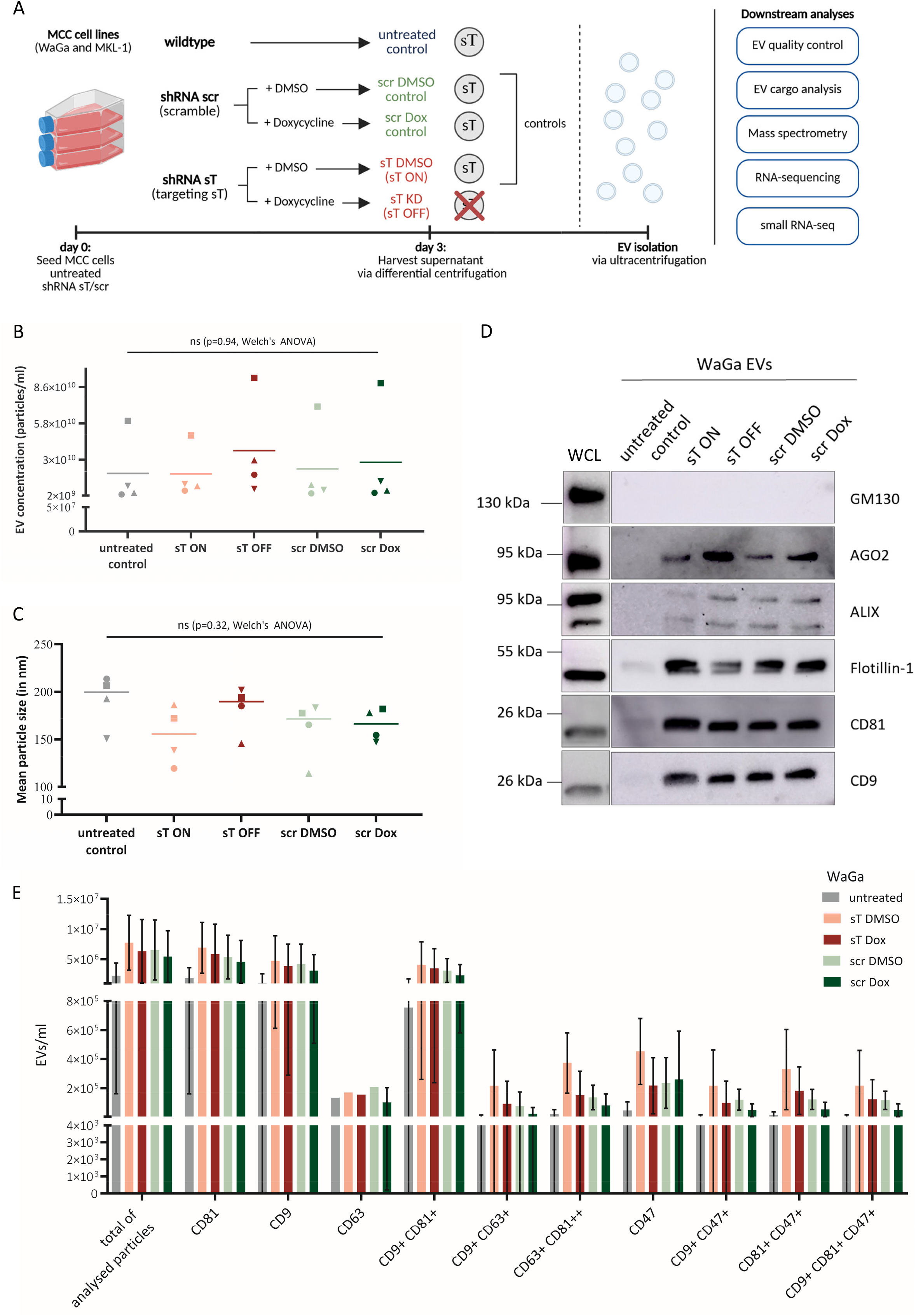
WaGa Merkel cell carcinoma cells release a heterogeneous extracellular vesicle (EV) population with characteristics of both exosomes and microvesicles. **(A)** Experimental workflow. Previously established WaGa and MKL-1 MCPyV-positive MCC cell lines carrying a doxycycline-inducible shRNA targeting sT (or a scrambled control) were used to modulate sT expression. EVs were isolated from conditioned medium of untreated cells and control or sT knockdown cultures by differential centrifugation followed by ultracentrifugation and subjected to quality control and cargo analyses. Knockdown of the viral small T antigen (sT) was induced by doxycycline treatment (sT OFF). DMSO-treated shRNA sT cells (sT ON), shRNA scramble (scr) cells treated with either DMSO or doxycycline, and untreated wild-type WaGa cells served as controls and retained sT expression. **(B–C)** Nanoparticle tracking analysis (NTA) of EVs isolated from WaGa cells. (B) EV particle concentration (particles/mL) and (C) mean particle diameter (nm) were determined. Individual dots represent independent biological experiments, and horizontal lines indicate the mean of four independent experiments. Statistical significance was assessed using Welch’s ANOVA, as appropriate (ns, not significant). **(D)** Western blot analysis of EV-associated proteins (Ago2, ALIX, Flotillin-1, CD81, and CD9) and the cellular marker GM130 in EV preparations from WaGa cells. Whole-cell lysates (WCL) served as positive controls for cellular protein expression. **(E)** Imaging flow cytometry (IFCM) analysis of EV surface markers (CD81, CD63, CD9, and CD47) on EVs isolated from WaGa cells. "Total analyzed particles" refers to all EVs that passed quality control and were included in the analysis. Data are presented as mean ± SD (n = 4; n = 1 for CD63 measurements)

Nanoparticle tracking analysis (NTA) revealed particle concentrations ranging from 1.9 × 10¹⁰ to 3.8 × 10¹⁰ particles/ml (Figure 1B, supplementary Figure 1D). The isolated particles exhibited a size distribution between approximately 100 and 250 nm, indicating a heterogeneous EV population that includes both small EVs (exosomes) and larger microvesicles (Figure 1C, supplementary Figure 1E). No significant differences in EV particle concentration or size distribution were observed across all experimental conditions. Western blot analysis confirmed the presence of the canonical EV markers ALIX, Flotillin-1, CD81, and CD9 as well as the RNA-binding protein AGO2, in WaGa-derived EVs (Figure 1D). The Golgi matrix protein GM130, used as a negative marker for cellular contamination, was detected only in WaGa whole-cell lysates (WCL) and was absent from the EV preparations, supporting the purity of the isolated EV fraction. Comparable findings were obtained for EVs derived from the independent MCPyV-positive MCC cell line MKL-1 (Supplementary Figure S1A-C), indicating that these characteristics are reproducible across MCPyV-positive MCC cell lines. WaGa-derived EVs were predominantly positive for CD81 and CD9, whereas CD63-positive EVs were detected at lower abundance (Figure 1E). A substantial fraction of EVs also expressed CD47, an immune regulatory protein previously shown to be upregulated by the MCPyV small T antigen (sT) in MCC cells and implicated in immune evasion through inhibition of macrophage-mediated phagocytosis (20). Efficient doxycycline-induced depletion of sT was verified at both the protein and transcript levels by Western blot analysis and RT-qPCR, respectively, in WaGa cells, with comparable knockdown efficiencies observed in MKL-1 cells (Supplementary Figure S2). Despite efficient sT depletion, no detectable changes in EV-associated CD47 levels were observed (Figure 1E, supplementary Figure 1F). Similarly, the proportions of the CD81⁺/CD47⁺, CD9⁺/CD47⁺, and CD9⁺/CD81⁺/CD47⁺ EV subpopulations remained comparable across all experimental conditions.

Cryo-electron microscopy revealed predominantly spherical membrane-enclosed vesicles of varying sizes with clearly visible lipid bilayers, consistent with the size distribution determined by NTA (supplementary Figure S1A, B). Several vesicles contained electron-dense material or additional internal membrane structures. Based on their morphology, EVs were classified into four categories: single vesicles, double vesicles, multilayer vesicles, and vesicles containing electron-dense cargo. Single vesicles represented the predominant population (43%), followed by multilayer vesicles (37.5%), vesicles containing electron-dense cargo (25%), and double vesicles (6%). Comparable morphological characteristics were observed for EVs derived from the independent MCPyV-positive MCC cell line MKL-1 (Supplementary Figure S1A, B), indicating that these features are reproducible across MCPyV-positive MCC cell lines. The observed morphological heterogeneity is consistent with previous cryo-electron microscopy studies of extracellular vesicles (46–48).

These findings demonstrate that MCC cells release a heterogeneous population of extracellular vesicles comprising both small and large EVs.

### WaGa-derived EVs exhibit a distinct proteomic profile and express canonical EV-associated proteins

To characterize the protein cargo of WaGa-derived EVs, mass spectrometry was performed, identifying 608 proteins consistently detected across all biological replicates. Comparison of the WaGa-derived EV proteome with the Top 100 EV-associated proteins in the Vesiclepedia database (version 5.1, September 2023) (49, 50) identified 72 overlapping proteins, supporting the enrichment of canonical EV-associated proteins in the isolated EV preparations (Figure 2A). Identified proteins included well-established EV-associated proteins such as CD81, CD9, ADAM10, and Annexin A5. In contrast, several commonly reported EV proteins, including CD63, Flotillin-1, and TSG101, were not reliably identified by LC-MS/MS-based proteomics, although Flotillin-1 was readily detected by Western blot analysis (Figure 1C). To determine the relationship between the EV proteome and that of the parental cells, the identified EV proteins were compared with the proteome of WaGa cells. Approximately 75% of the EV proteins were also detected in the parental cells (Figure 2B), indicating that the EV protein cargo partially reflects the molecular composition of the donor cells. Notably, 148 proteins were detected exclusively in EVs, whereas numerous cellular proteins were absent from the EV fraction, demonstrating that the EV proteome differs substantially from that of the parental cells. To further characterize the molecular composition of WaGa-derived EVs and the parental cells, pathway enrichment analysis was performed using the DAVID functional annotation tool. Reactome pathway enrichment analysis identified proteins associated with transcriptional regulation, DNA repair, antigen processing and presentation, and MAPK and AKT signaling pathways (Figure 2C, D). Comparable proteomic characteristics were observed for MKL-1-derived EVs (supplementary Figure S3). Specifically, 82 of the Top 100 EV-associated proteins listed in the Vesiclepedia database were identified, and comparison of the EV and parental cell proteomes revealed a similar pattern of shared and EV-associated proteins, supporting the reproducibility of these findings in an independent MCPyV-positive MCC cell line. Together, these results show that MCC-derived EVs possess a distinct proteomic composition that shares core EV-associated proteins with established EV databases while differing substantially from the proteome of their parental cells.

**Figure 2.**
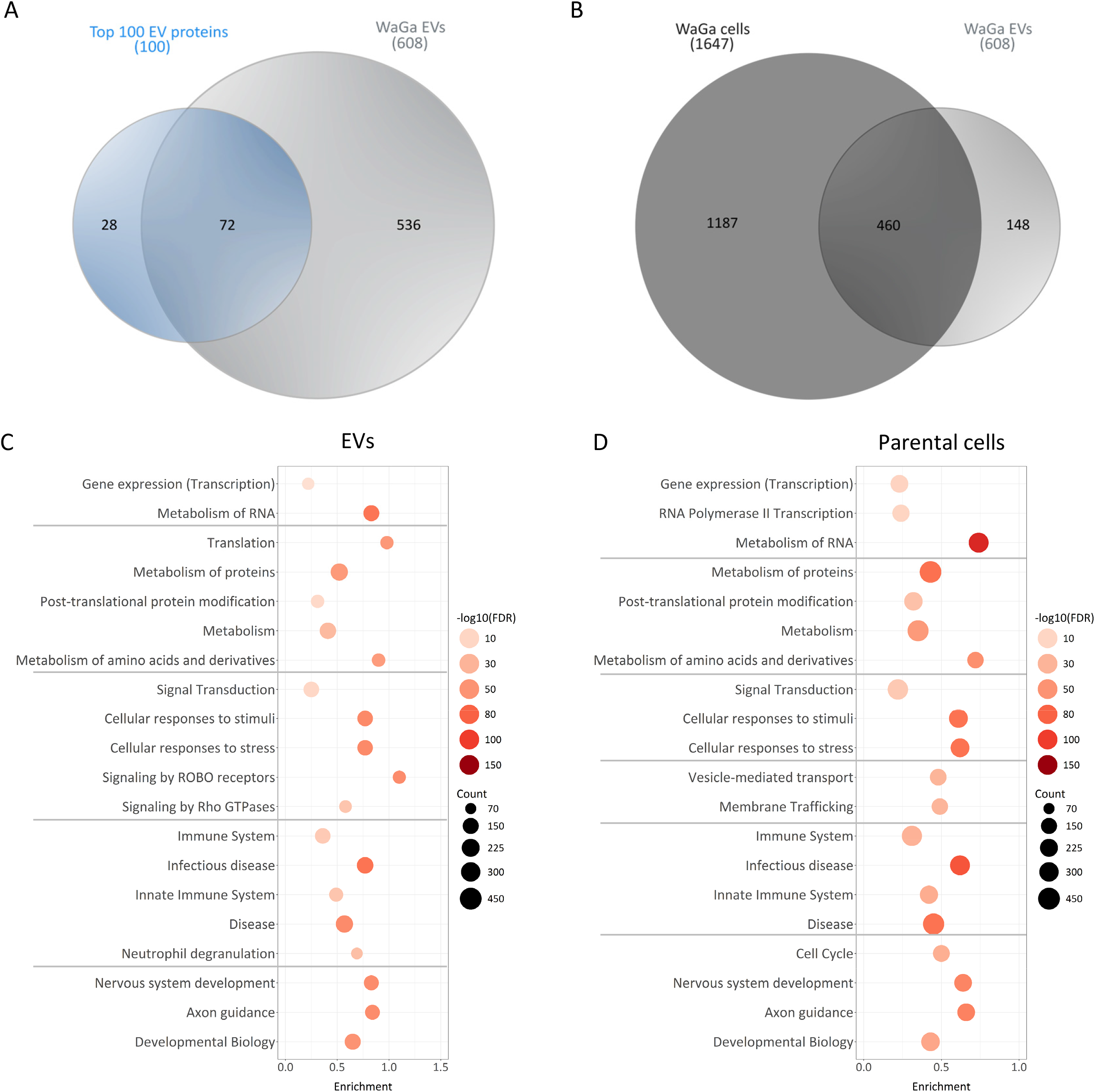
WaGa-derived EVs exhibit a distinct proteomic profile and express canonical EV marker proteins. **(A)** Venn diagram comparing the Top 100 extracellular vesicle (EV) proteins listed in the Vesiclepedia database (Version 5.1, September 2023 release) with the proteins identified in EVs isolated from untreated WaGa cells. **(B)** Venn diagram comparing the proteome of untreated WaGa whole-cell lysates with the proteome of EVs isolated from untreated WaGa cells. **(C–D)** Reactome pathway enrichment analysis was performed using the DAVID functional annotation tool. (C) Analysis of all proteins identified in EVs isolated from untreated WaGa cells. (D) Analysis of all proteins identified in untreated WaGa whole-cell lysates. Among all significantly enriched Reactome pathways (false discovery rate, FDR ≤ 0.05), the 20 pathways with the highest gene counts were selected and grouped according to their biological function. Circle size represents the number of proteins assigned to each pathway, whereas color intensity indicates the level of statistical significance (FDR ≤ 0.05).

### MCPyV MCC cell-derived EVs display selective enrichment of specific RNA species relative to their parental cells

To characterize the RNA cargo of WaGa and MKL-1-derived EVs and compare it with that of the parental cells, total RNA sequencing was performed on EVs and corresponding WaGa cells. Protein-coding transcripts represented the majority of annotated reads in both sample types, accounting for approximately 93% of reads in WaGa and MKL-1 cells and 88% and 85% in EVs from WaGa and MKL-1 cells respectively (Supplementary Figure S4A). Compared with the parental cells, EVs exhibited a higher relative abundance of long non-coding RNAs (lncRNAs) and pseudogenes, indicating differences in RNA cargo composition between EVs and their cells of origin. Initial characterization of EV-associated RNA revealed marked quality differences compared with RNA from the corresponding parental cells (Supplementary Figure S4B). Electrophoretic analysis showed that EV RNA was enriched for shorter RNA species and lacked the prominent high-molecular-weight RNA bands observed in parental-cell RNA. Consistent with this distinct RNA composition, RNA-sequencing reads from WaGa EVs showed slightly lower overall alignment rates (60.7–68.1%) and a lower proportion of reads assigned to protein-coding genes (82.4–87.9%) compared with parental-cell RNA (71.0% and 92.9%, respectively). Notably, EV-associated reads displayed a pronounced shift in their genomic distribution, with a lower proportion mapping to exons (15.1–28.5% versus 59.8% in parental cells) and a higher proportion mapping to introns (44.6–52.7% versus 34.0%). Similar results were obtained for MKL-1 RNA sequencing from EVs and parental cells (Supplementary Figure S4B-C). To further compare the RNA cargo of EVs and parental cells, differential abundance analysis was performed. Approximately 26,000 transcripts showed a higher relative abundance in EVs, whereas approximately 2,800 transcripts exhibited a lower relative abundance compared with the parental cells (Figure 3B), demonstrating that the RNA composition of WaGa-derived EVs differs substantially from that of their cells of origin. Comparable differences in RNA cargo composition were observed in EVs isolated from the independent MCPyV-positive MCC cell line MKL-1 (Supplementary Figure S5A), indicating that these findings are reproducible across MCPyV-positive MCC cell lines.

**Figure 3:**
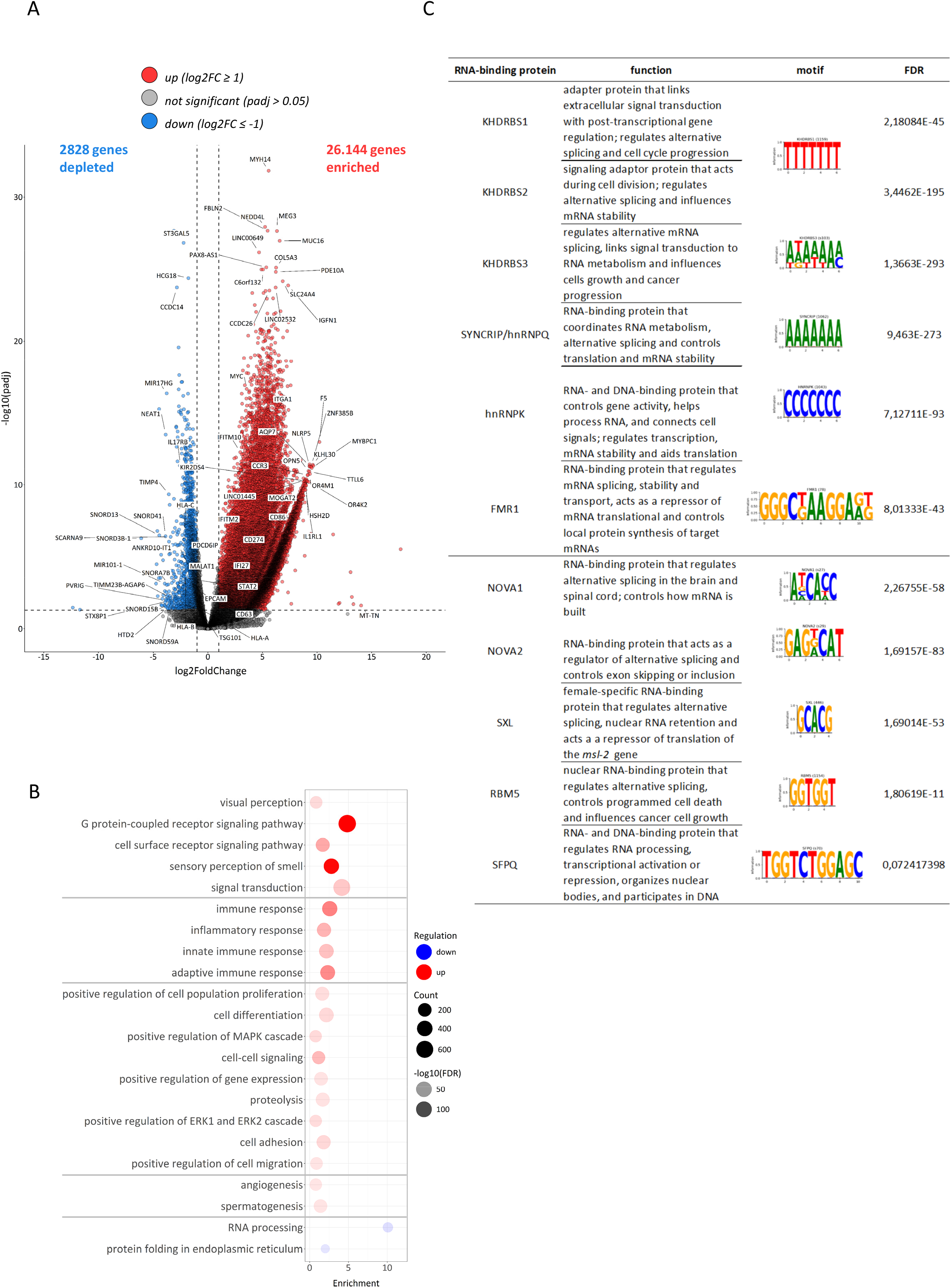
WaGa-derived EV RNA cargo is enriched for transcripts associated with immune signaling, cell communication, and RNA processing. **(A)** Volcano plot showing differentially expressed genes (DEGs) in EVs isolated from untreated WaGa cells compared with parental WaGa cells. Genes with a log2 fold change > 1 or < –1 and an adjusted *P* value (padj.) < 0.05 were considered significantly upregulated or downregulated, respectively. **(B)** Gene Ontology (GO) Biological Process enrichment analysis of significantly differentially expressed genes identified in (B) was performed using the DAVID functional annotation tool. Among all significantly enriched GO terms (FDR ≤ 0.05), the 20 terms with the highest gene counts were selected and grouped according to their biological function. Circle size represents the number of genes associated with each GO term, whereas color indicates the direction of regulation (red, enriched among upregulated genes; blue, enriched among downregulated genes). Color intensity reflects the level of statistical significance (FDR ≤ 0.05). **(C)** RNA-binding protein (RBP) motif enrichment analysis of transcripts enriched in untreated WaGa EVs using the ATtRACT database. Significantly enriched motifs (FDR < 0.1) are shown together with the corresponding RBP and FDR.

To investigate the biological functions associated with transcripts exhibiting higher relative abundance in EVs, Gene Ontology (GO) biological process enrichment analysis was performed using the DAVID functional annotation tool. Transcripts with higher relative abundance in EVs were associated with biological processes including cell signaling, immune response, cell proliferation, and angiogenesis (Figure 3B). In contrast, transcripts with lower relative abundance in EVs were primarily associated with RNA processing and protein folding, reflecting differences in transcript representation between EVs and their parental cells.

To determine whether EV-associated transcripts contain enriched RNA-binding protein (RBP) recognition motifs, Analysis of Motif Enrichment (AME) was performed using the ATtRACT database. Several significantly enriched sequence motifs and their corresponding RBPs were identified (Figure 3C). These included T-rich motifs recognized by KHDRBS1/2, A-rich motifs bound by KHDRBS3 and SYNCRIP (hnRNP Q), C-rich motifs associated with hnRNP K, and G-rich motifs recognized by FMR1 and RBM5. In addition, CA-rich and GC-rich core motifs corresponding to binding sites for NOVA1/2 and SXL, respectively, were significantly enriched among EV-associated transcripts. Similar RBP-binding motif enrichment patterns were identified in MKL-1-derived EVs (Supplementary Figure S5), indicating that similar RBP-binding motifs are represented in EV-associated transcripts from both MCC cell lines.

These findings demonstrate that WaGa-derived EVs contain an RNA cargo composition that differs from that of their parental cells and is characterized by enrichment of transcripts associated with signaling- and immune-related biological processes together with distinct RNA-binding protein recognition motifs.

### WaGa-derived EVs display a distinct miRNA cargo compared with their parental cells

To comprehensively characterize the small RNA cargo of WaGa-derived EVs, parental cells and EVs were subjected to small RNA sequencing. Sequencing data were analyzed using the exceRpt pipeline, which is optimized for extracellular RNA analysis. Approximately 98% of reads obtained from WaGa cells mapped to the human genome, whereas approximately 73% of reads from WaGa-derived EVs could be mapped. In parental cells, 59% of mapped reads were annotated as miRNAs and 32% as long RNAs, whereas tRNAs and piRNAs accounted for approximately 2% and 0.3%, respectively (Figure 4A, results doe MKL-1 cells are shown in Suppl. Figure S6A). In WaGa-derived EVs, 28% of mapped reads were annotated as miRNAs and 10% as long RNAs, while the proportion of tRNA-derived reads increased to approximately 29% and piRNA-derived reads accounted for 0.65%. These findings demonstrate marked differences in small RNA composition between WaGa-derived EVs and their parental cells.

**Figure 4.**
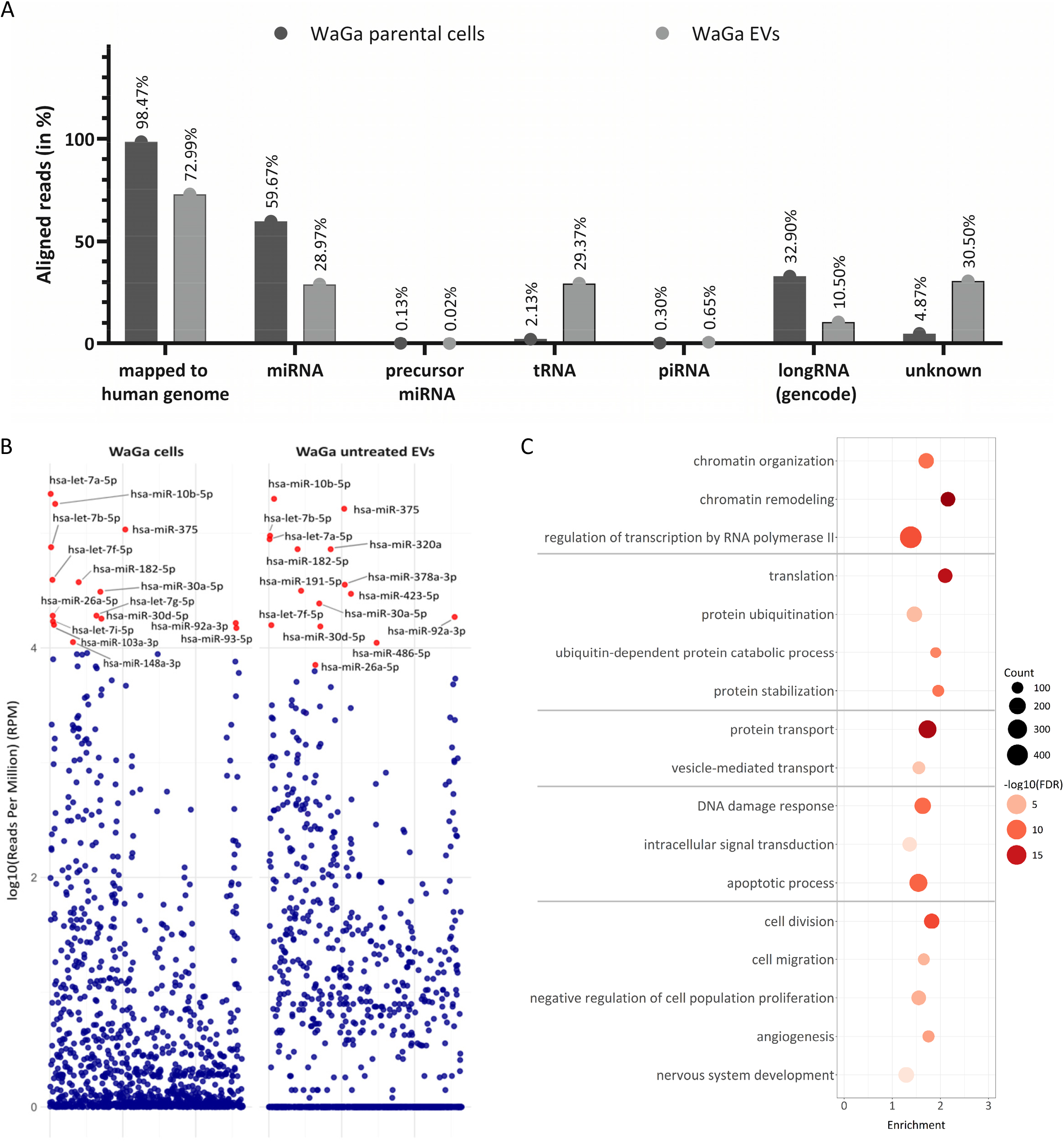
WaGa-derived EVs are enriched in miRNAs predicted to target cancer-related biological processes. **(A)** Small RNA sequencing reads were aligned to the human genome, and the abundance of individual small RNA biotypes identified in WaGa-derived EVs was determined. Numbers indicate the total number of aligned reads and the relative contribution of each small RNA class to all annotated reads (%). **(B)** Manhattan plots showing normalized miRNA abundance (log10 reads per million; RPM) in parental WaGa cells (left) and untreated WaGa-derived EVs (right). Red dots indicate the 15 most abundant miRNAs (by RPM) in each sample type. Each dot represents one miRNA; data are derived from two independent biological replicates (n = 2). **(C)** Gene Ontology (GO) Biological Process enrichment analysis of the predicted target genes of the 15 most abundant miRNAs identified in (B) was performed using the DAVID functional annotation tool. Among all significantly enriched GO terms (FDR ≤ 0.05), those with the highest gene counts were selected and grouped according to their biological function. Circle size represents the number of target genes associated with each GO term, whereas color intensity reflects the level of statistical significance (FDR ≤ 0.05).

The miRNA cargo of WaGa-derived EVs was subsequently analyzed in greater detail. Among the 15 most abundant miRNAs detected in WaGa-derived EVs were miR-375, miR-10b-5p, miR-182-5p, and miR-30a-5p (Figure 4B). Comparison with the parental cells showed that most miRNAs represented among the 15 most abundant miRNAs exhibited similar relative abundance in EVs and cells. However, miR-378a-3p, miR-423-5p, and miR-485-5p were overrepresented in WaGa-derived EVs relative to the parental cells. These findings indicate that the miRNA cargo of WaGa-derived EVs largely reflects highly abundant cellular miRNAs, while a subset of miRNAs shows increased representation within the EV fraction.

Analysis of MKL-1-derived EVs revealed a partly distinct miRNA profile (supplementary Figure S6B). In contrast to WaGa-derived EVs, miR-378a-3p, miR-423-5p, and miR-485-5p were not among the 15 most abundant miRNAs in MKL-1 EVs. Instead, miR-25-3p, miR-92b-3p, and miR-98-5p were represented among the most abundant miRNAs in MKL-1-derived EVs, demonstrating that the highly abundant EV-associated miRNA repertoire differs between the two MCPyV-positive MCC cell lines.

To explore the potential biological functions of EV-associated miRNAs, experimentally validated target genes were retrieved from the miRTarBase database and subjected to Gene Ontology biological process enrichment analysis. Target genes were associated with biological processes including gene regulation, translation, DNA damage response, cellular regulation, and apoptosis (Figure 4C). Despite differences in the repertoire of highly abundant EV-associated miRNAs between WaGa and MKL-1 cells, GO enrichment analysis identified largely similar biological processes in both cell lines (Supplementary Figure S6C), suggesting that distinct EV miRNA profiles converge on common functional pathways.

Together, these findings show that the miRNA cargo of MCC-derived EVs is composed predominantly of abundant cellular miRNAs but also includes individual miRNAs that are overrepresented relative to the parental cells. Moreover, the repertoire of highly abundant EV-associated miRNAs differs between WaGa and MKL-1 cells, indicating cell-line-specific variation in EV miRNA composition.

### Network analysis identifies highly connected target genes of abundant EV-associated miRNAs

To investigate the regulatory network associated with abundant EV miRNAs, experimentally validated target genes of the 15 most abundant miRNAs identified in WaGa-derived EVs were retrieved from miRTarBase and used to construct a miRNA-target interaction network (Figure 5A). The resulting network comprised 196 target genes, illustrating the extensive regulatory interactions between abundant EV-associated miRNAs and their downstream targets.

**Figure 5.**
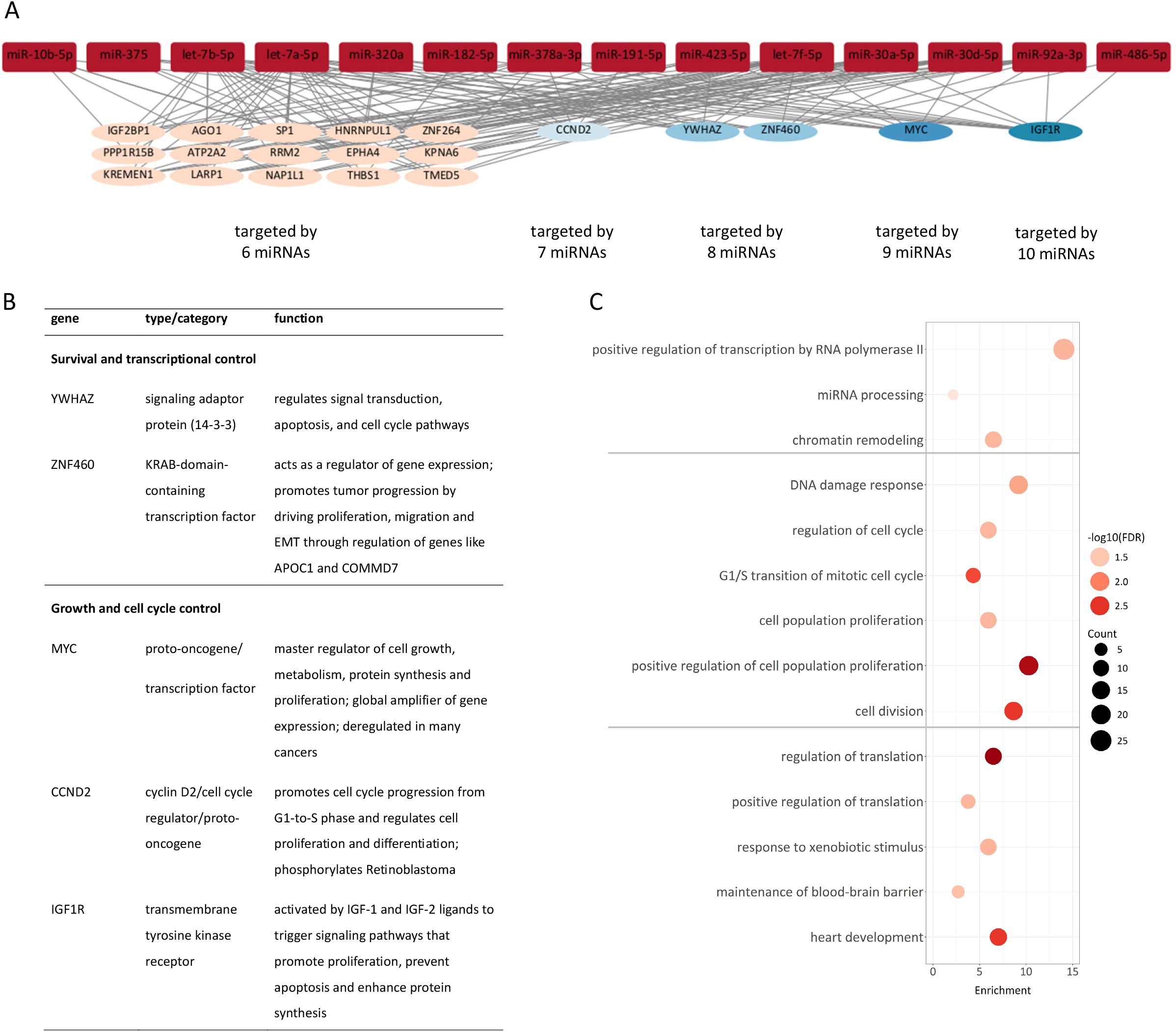
Network analysis identifies genes targeted by multiple EV-associated miRNAs. **(A)** Network analysis of experimentally validated target genes (miRTarBase) of the 10 most enriched EV-associated miRNAs. miRNAs are shown as dark red nodes, whereas target genes are color-coded according to the number of miRNAs predicted to target them. Edges indicate experimentally validated miRNA–target gene interactions. **(B)** Genes targeted by multiple miRNAs, identified in the network analysis shown in panel A, together with their annotated biological functions. **(C)** Gene Ontology (GO) biological process enrichment analysis of the multi-targeted genes was performed using DAVID. Among all significantly enriched GO terms (FDR ≤ 0.05), those with the highest gene counts were selected and grouped according to their biological function. Circle size corresponds to the number of associated genes, and color intensity indicates the significance of enrichment (FDR ≤ 0.05).

To identify potential regulatory hubs, target genes were grouped according to the number of distinct miRNAs targeting each gene (Figure 5A). While the majority of genes were targeted by relatively few miRNAs, a subset of genes exhibited substantially higher connectivity within the network. Notably, IGF1R and MYC represented the most highly connected nodes and were targeted by 10 and 9 of the 15 most abundant EV-associated miRNAs, respectively. Additional highly connected target genes included CCND2, YWHAZ, THBS1, AGO1, KPNA6, and SP1, indicating that multiple abundant EV-associated miRNAs converge on a limited number of shared target genes (Figure 5A, B).

To assess the biological functions associated with these highly connected target genes, Gene Ontology (GO) biological process enrichment analysis was performed. The analysis identified significant enrichment of processes related to transcriptional regulation, miRNA processing, DNA damage response, cell cycle regulation, cell proliferation, cell division, and regulation of translation (Figure 5C).

Comparable network architecture was observed for MKL-1-derived EVs (Supplementary Figure S7). Although the identity of the most abundant EV-associated miRNAs and their highest connected target genes differed from those identified in WaGa-derived EVs, the network similarly contained a subset of highly connected regulatory hubs targeted by multiple miRNAs. Moreover, GO enrichment analysis revealed largely overlapping biological processes, including transcriptional regulation, miRNA processing, cell cycle regulation, apoptosis, translation, and cellular stress responses, indicating that distinct EV-associated miRNA repertoires in MCPyV-positive MCC cell lines converge on similar functional pathways.

These findings support that abundant EV-associated miRNAs form complex regulatory networks characterized by highly connected target genes and conserved functional signatures despite cell line-specific differences in individual miRNAs.

### Knockdown of the viral small T antigen has limited effects on the RNA cargo of WaGa-or MKL-1 derived EVs

To investigate whether the viral small T antigen (sT) influences the RNA cargo of EVs, small RNA sequencing and total RNA sequencing were performed on EVs isolated from WaGa and MKL-1 cells cultured under sT knockdown (sT Dox; sT OFF) or control (sT DMSO; sT ON) conditions (Figure 6, Supplementary Figure S8-S11). Efficient sT depletion was confirmed by Western blotting and target-specific RT-qPCR before EV isolation (supplementary Figure S2). Comparison of the EV-associated miRNA profiles revealed no significant differences between sT knockdown and control cells (Supplementary Figure S8), indicating that depletion of sT does not substantially alter the miRNA composition of WaGa-derived EVs.

**Figure 6.**
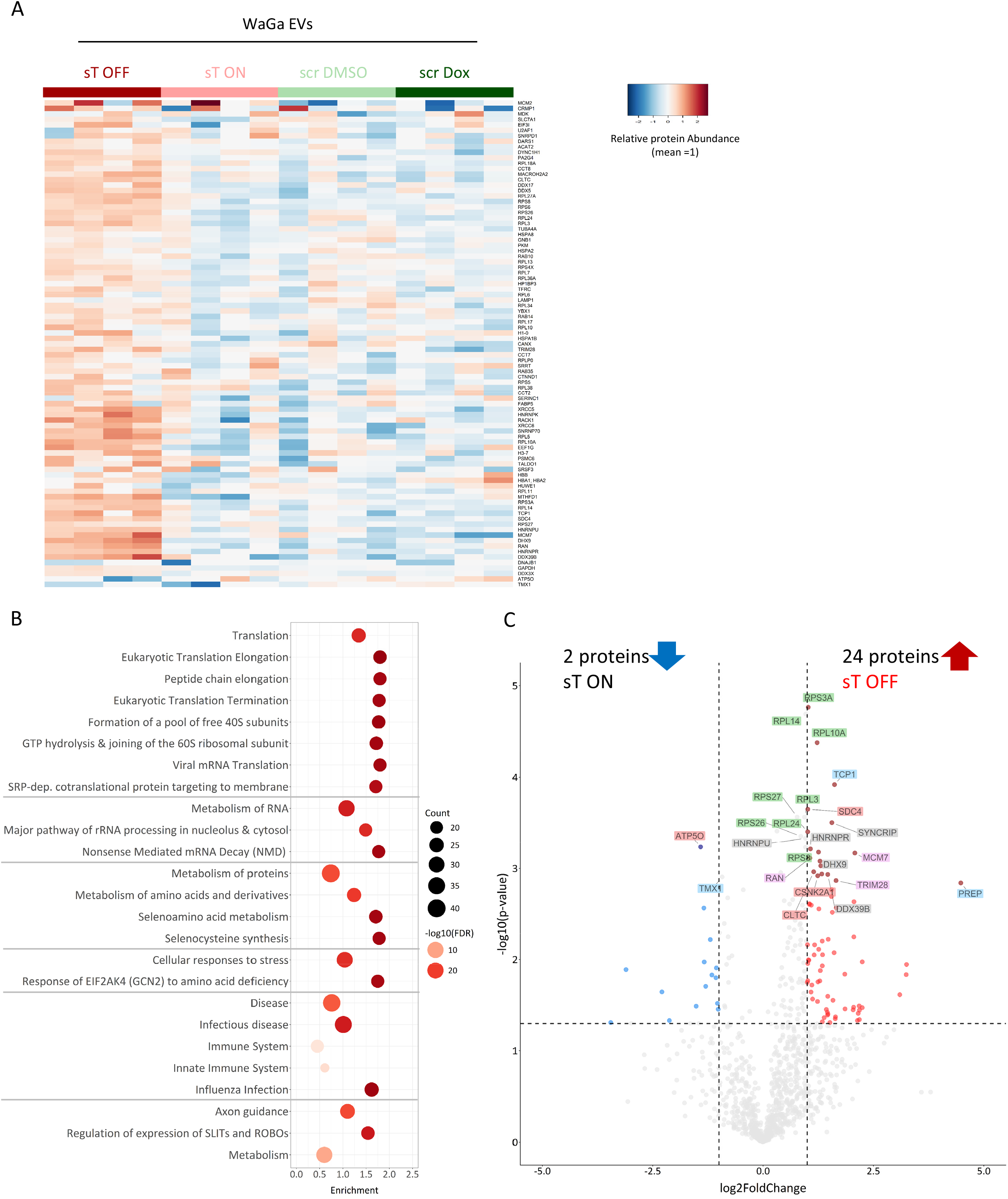
Knockdown of the viral small T antigen (sT) reshapes the proteomic cargo of WaGa-derived extracellular vesicles. **(A)** Heat map showing the relative abundance of proteins identified by mass spectrometry in EVs isolated from WaGa shRNA sT and shRNA scramble (scr) cells treated with DMSO or doxycycline. Color intensity indicates relative protein abundance. **(B)** Reactome pathway enrichment analysis of proteins enriched in EVs derived from doxycycline-treated WaGa shRNA sT cells (sT Dox) was performed using the DAVID functional annotation tool. Among all significantly enriched Reactome pathways (FDR ≤ 0.05), the pathways with the highest gene counts were selected. Circle size represents the number of proteins assigned to each pathway, whereas color intensity reflects the level of statistical significance (FDR ≤ 0.05). **(C)** Volcano plot of proteomic data showing differentially abundant proteins in EVs derived from doxycycline-treated WaGa shRNA sT cells (sT OFF) compared with the corresponding doxycycline-treated shRNA scramble control (scr Dox; sT ON). Proteins with a log2 fold change > 1 or < −1 and an adjusted *P* value (padj.) < 0.05 were considered significantly increased or decreased, respectively. Ribosomal /translational proteins are shown in green; RNA processing proteins are shown in grey; Protein folding/quality control proteins are shown in blue; Genome maintenance proteins are shown in purple; signaling/cellular organization proteins are shown in red.

Hierarchical clustering demonstrated considerable variability among biological replicates but also revealed reproducible differences between control and sT knockdown EVs (supplementary Figure S10A). Differential abundance analysis identified 15 transcripts with significantly higher relative abundance following sT knockdown, whereas no transcripts exhibited significantly lower abundance (supplementary Figure S10B). Among the transcripts with increased abundance were LIG1 and ARGLU1, genes previously implicated in DNA damage responses, cell survival, and tumor progression. Collectively, these findings indicate that knockdown of sT has only a limited impact on the RNA cargo of WaGa-derived EVs, with no detectable changes in the EV-associated miRNA profile and only a small number of mRNAs exhibiting altered abundance.

### Knockdown of the viral small T antigen is associated with changes in the proteomic cargo of WaGa-derived EVs

To determine whether sT also influences the protein cargo of extracellular vesicles, differential quantitative mass spectrometry-based proteomics was performed on EVs isolated from sT knockdown (sT Dox) and control (scr Dox) WaGa cells. Hierarchical clustering of the identified proteins revealed considerable variability among biological replicates but also identified a subset of proteins exhibiting consistently altered abundance following sT knockdown (Figure 6A). Reactome pathway enrichment analysis of the differentially abundant proteins identified pathways related to cellular stress responses, translation, RNA metabolism, and the immune system (Figure 6B).

Differential abundance analysis identified 26 proteins whose abundance was significantly altered following sT knockdown, comprising 24 proteins with higher abundance and 2 proteins with lower abundance in EVs (Figure 6C). Proteins exhibiting increased abundance included several ribosomal proteins (RPS3A, RPL14, and RPL10A), the RNA-binding proteins hnRNP U and SYNCRIP (hnRNP Q), as well as TRIM28. In contrast, only ATP5O and TMX1 showed significantly lower abundance following sT knockdown.

Analysis of EVs isolated from the independent MCPyV-positive MCC cell line MKL-1 identified only two differentially abundant proteins following sT knockdown (Supplementary Figure S11), indicating that the proteomic response to sT depletion differs between the two MCPyV-positive MCC cell lines. Collectively, these findings demonstrate that depletion of sT is associated with distinct, but relatively limited, changes in the proteomic cargo of WaGa-derived EVs.

## Discussion

Although the cell-intrinsic functions of the MCPyV oncoproteins have been extensively studied, comparatively little is known about how MCPyV-positive MCC cells communicate with the tumor microenvironment through extracellular vesicles. In this study, we provide the first integrated multi-omics characterization of the protein and RNA cargo of EVs released by MCPyV-positive MCC cells and investigate whether the viral oncoprotein sT contributes to shaping this cargo. A central finding of this study is that, although EV cargo composition differs between the two MCPyV-positive MCC cell lines, their protein and RNA cargoes converge on remarkably similar biological functions, suggesting conservation of functional EV signatures despite molecular heterogeneity.

Our analyses demonstrate that MCPyV-positive MCC cells release a heterogeneous population of EVs displaying characteristics consistent with both small and large EVs. This was supported by nanoparticle tracking analysis, imaging flow cytometry, transmission electron microscopy, and the detection of canonical EV-associated proteins. Importantly, comparable physicochemical and molecular characteristics were observed in two independent MCPyV-positive MCC cell lines, WaGa and MKL-1, indicating that these findings are reproducible and not restricted to a single cellular model.

Proteomic profiling revealed substantial overlap between MCC-derived EV proteins and the Top 100 EV-associated proteins listed in Vesiclepedia, confirming the successful isolation of bona fide EVs. Nevertheless, mass spectrometry failed to detect several canonical EV markers, including ALIX, Flotillin-1 and CD63, which were readily detected by Western blotting and imaging flow cytometry. Similar discrepancies have been reported previously and likely reflect differences in the analytical sensitivity and dynamic range of antibody-based and mass spectrometry-based approaches rather than the absence of these proteins from the EV preparations. Together, these observations highlight the importance of combining complementary methods for EV characterization, in agreement with current MISEV recommendations.

Although a substantial proportion of the EV proteome overlapped with that of the parental cells, approximately one quarter of the detected proteins were identified exclusively in EVs, indicating that the protein cargo is not merely a random representation of the cellular proteome. Instead, these data are consistent with the concept that EVs possess a distinct protein composition and that specific proteins become relatively enriched during EV biogenesis. Interestingly, histone proteins represented some of the most abundant proteins preferentially detected in EVs. Histones have previously been identified in EVs from several tumor entities and have been implicated in chromatin-associated signaling, inflammatory responses, and modulation of recipient cells. Whether histone-enriched EVs contribute to MCC biology remains to be determined.

RNA sequencing further demonstrated that MCC-derived EVs contain a distinct RNA cargo compared with their parental cells. Besides protein-coding transcripts, EVs contained a broad spectrum of RNA species and exhibited enrichment of transcripts associated with RNA metabolism, transcriptional regulation, intracellular transport and signal transduction. Motif enrichment analysis identified multiple RNA-binding protein (RBP) recognition motifs among EV-enriched transcripts. Several of the corresponding RBPs, including members of the heterogeneous nuclear ribonucleoprotein (hnRNP) family, have previously been implicated in RNA trafficking and EV cargo selection. While our data do not demonstrate direct involvement of these RBPs in MCC, they are consistent with previous studies suggesting that RNA incorporation into EVs is influenced, at least in part, by sequence-dependent recognition mechanisms.

The small RNA cargo similarly differed from that of the parental cells. Although the most abundant EV-associated miRNAs varied between WaGa and MKL-1 cells, both cell lines displayed remarkably similar functional enrichment profiles. Target gene analyses consistently identified biological processes involved in transcriptional regulation, cell cycle control, apoptosis, DNA damage responses and intracellular signaling. Likewise, network analysis demonstrated that despite differences in the identity of the predominant miRNAs and their highest connected target genes, the overall regulatory networks converged on similar biological functions. These observations suggest that distinct EV-associated miRNA repertoires may regulate overlapping cellular pathways, thereby preserving functional output despite differences in individual miRNA composition. Interestingly, a similar pattern emerged from both the proteomic and transcriptomic analyses, indicating that molecular differences between MCC-derived EVs converge on conserved functional programs across independent cell lines.

Among the most abundant EV-associated miRNAs, miR-375 was consistently detected in both MCC cell lines. Previous studies demonstrated that EV-mediated transfer of miR-375 promotes polarization of cancer-associated fibroblasts and contributes to the establishment of a tumor-supportive microenvironment in MCC. Our findings extend these observations by demonstrating that miR-375 is embedded within a broader EV-associated miRNA network predicted to regulate multiple pathways involved in tumor progression. Although these analyses do not establish functional activity of the transferred miRNAs, they provide a framework for future studies investigating the contribution of MCC-derived EVs to microenvironmental remodeling.

A major objective of this study was to determine whether the viral oncoprotein sT influences EV cargo composition. Surprisingly, inducible depletion of sT resulted only in relatively modest alterations of the EV cargo. Proteomic analysis identified a limited number of proteins with increased relative abundance following sT knockdown, many of which were associated with ribosome biogenesis, RNA metabolism or RNA-binding functions. Likewise, only fifteen transcripts exhibited significantly higher relative abundance in EVs after sT depletion, whereas no transcripts showed significantly reduced abundance. In contrast, the overall EV-associated miRNA landscape remained largely unchanged, with only a small subset of miRNAs displaying altered abundance following sT knockdown. These findings indicate that the global composition of the EV cargo is largely maintained despite loss of sT.

Interestingly, several proteins exhibiting increased abundance following sT depletion, including hnRNP family members, have previously been linked to RNA processing and EV biology. Whether these changes represent compensatory responses to altered cellular homeostasis or reflect indirect consequences of sT depletion remains unknown. Likewise, the enrichment of transcripts associated with transcriptional regulation and RNA processing following sT knockdown may indicate subtle alterations in RNA trafficking rather than large-scale remodeling of EV cargo composition. Functional experiments will be required to determine whether these molecules influence communication between MCC cells and their microenvironment.

Our study has several limitations. First, all analyses were performed using established MCC cell lines cultured in vitro, which cannot fully recapitulate the complexity of the tumor microenvironment. Second, the functional consequences of EV-mediated transfer of the identified proteins and RNAs were not addressed experimentally. Finally, although two independent MCPyV-positive MCC cell lines were analyzed, future studies should include primary patient-derived EVs to determine the clinical relevance of the identified cargo signatures.

In conclusion, we provide the first comprehensive multi-omics characterization of extracellular vesicles released by MCPyV-positive MCC cells. We demonstrate that MCC-derived EVs possess distinct protein, mRNA and miRNA cargoes compared with their parental cells. Despite substantial differences in the molecular composition of EV cargo between WaGa and MKL-1 cells, functional enrichment analyses consistently identified highly similar biological pathways, indicating convergence on conserved EV-associated functions. Furthermore, depletion of the viral oncoprotein sT induced only modest changes in EV cargo composition. Furthermore, we show that depletion of the viral oncoprotein sT induces only modest changes in EV cargo composition, suggesting that the overall molecular architecture of MCC-derived EVs is largely maintained independently of sT expression. Together, these findings establish a resource for future investigations into EV-mediated communication in MCC and provide a foundation for exploring the contribution of EV cargo to tumor–microenvironment interactions and disease progression.

## Data Availability

All RNA sequencing and small RNA sequencing data generated in this study have been deposited in the Gene Expression Omnibus (GEO) under the SuperSeries accession number GSE343032 (total RNA sequencing) and GSE343035 (small RNA sequencing). The RNA sequencing dataset (GSE343032) comprises 40 samples, including parental cells, wild-type EVs, and EVs derived from shRNA sT or shRNA scramble cells treated with DMSO or doxycycline from both WaGa and MKL-1 cell lines. The small RNA sequencing dataset (GSE343035) comprises 34 samples covering the same experimental conditions.

## Acknowledgements

We thank Kerstin Reumann for expert technical assistance.

The project was funded by the Deutsche Forschungsgemeinschaft (DFG, German Research Foundation) - GRK2771 – project no. 453548970

## Author contribution statement

Conceptualization: N.F.

Methodology: U.W., B.S., A.S.S., P.B, N.F.

Investigation: U.W., J.H., C.S.

Formal analysis: U.W., B.S., J. H., N.F.

Resources: N.F.

Writing – original draft: U.W, N.F.

Funding acquisition: N.F.

All authors critically reviewed the manuscript, approved the final version, and agree to be accountable for all aspects of the work.

## Supplementary Information

- ***Supplementary Figures S1-S11***
- ***Supplementary Tables S1-S3***

***Table S1**: Top 100 most abundant proteins identified in extracellular vesicles released by WaGa and MKL-1 cells*.

***Table S2**: Reactome pathway enrichment analysis of the WaGa-derived extracellular vesicle proteome shown in Figure 2C*.

***Table S3:** Twenty-five most abundant miRNAs identified in WaGa and MKL-1 cells and their corresponding extracellular vesicles*.

## Figure legends Supplementary Figures

**Figure S1: Cryo-electron microscopy reveals morphologically heterogeneous WaGa-derived and MKL-1-derived extracellular vesicles with a visible lipid bilayer.**

**(A)** Representative cryo-electron microscopy (cryo-EM) images of EVs isolated from untreated WaGa cells. Images illustrate representative EV morphologies corresponding to the categories shown in (B).

**(B)** Quantification of EV morphologies identified by cryo-EM. A total of 16 EVs were imaged and classified according to their morphology. Categories were not mutually exclusive.

**(C)** Western blot analysis of EV-associated proteins (Ago2, ALIX, Flotillin-1, CD81, and CD9) and the cellular marker GM130 in EV preparations from MKL-1 cells. Whole-cell lysates (WCL) served as positive controls for cellular protein expression.

**(D–E)** Nanoparticle tracking analysis (NTA) of EVs isolated from MKL-1 cells. (D) EV particle concentration (particles/mL) and (E) mean particle diameter (nm) were determined. Individual dots represent independent biological experiments, and horizontal lines indicate the mean of four independent experiments. Statistical significance was assessed using Welch’s ANOVA, as appropriate (ns, not significant).

**(F)** Imaging flow cytometry (IFCM) analysis of EV surface markers (CD81, CD63, CD9, and CD47) on EVs isolated from MKL-1 cells. "Total analyzed particles" refers to all EVs that passed quality control and were included in the analysis. Data are presented as mean ± SD.

**Figure S2: Validation of doxycycline-induced sT knockdown in WaGa cells.**

**(A)** Representative Western blot showing sT protein expression in WaGa cells carrying either the inducible shRNA sT or shRNA scramble (scr) construct following treatment with DMSO or doxycycline (Dox). β-actin served as a loading control. **(B)** Quantification of relative sT transcript levels following DMSO or doxycycline treatment. Expression levels were normalized to ACTB (β-actin) and are shown relative to the DMSO-treated control (set to 1). Data are presented as mean ± SD. (C) RT-qPCR analysis of relative sT transcript levels in WaGa cells. Expression levels were normalized to GAPDH and are shown relative to the DMSO-treated control (set to 1). Data are presented as mean ± SD. **(D)** RT-qPCR analysis of relative sT transcript levels in MKL-1 cells. Expression levels were normalized to GAPDH and are shown relative to the DMSO-treated control (set to 1). Data are presented as mean ± SD.

**Figure S3: Proteomic characterization of untreated MKL-1-derived EVs.**

**(A)** Venn diagram comparing the Top 100 EV-associated proteins listed in the Vesiclepedia database (Version 5.1, September 2023) with the proteins identified in MKL-1-derived EVs.

**(B)** Venn diagram comparing the proteins identified in MKL-1 cells with those identified in MKL-1-derived EVs.

**(C–D)** Reactome pathway enrichment analysis was performed using the DAVID functional annotation tool. (C) Analysis of all proteins identified in EVs isolated from untreated MKL-1 cells. (D) Analysis of all proteins identified in untreated MKL-1 whole-cell lysates. Among all significantly enriched Reactome pathways (false discovery rate, FDR ≤ 0.05), the 20 pathways with the highest gene counts were selected and grouped according to their biological function. Circle size represents the number of proteins assigned to each pathway, whereas color intensity indicates the level of statistical significance (FDR ≤ 0.05).

**Figure S4: Characterization of RNA composition and sequencing features of WaGa- and MKL-1-derived extracellular vesicles**

**(A)** Distribution of RNA biotypes in parental cells and extracellular vesicles. Relative proportions (%) of reads assigned to different RNA biotypes in total RNA-sequencing data from WaGa and MKL-1 parental cells and their corresponding extracellular vesicles (EVs). RNA biotypes include protein-coding RNA, long non-coding RNA (lncRNA), pseudogene-derived RNA, small RNA, ribosomal RNA (rRNA), transfer RNA (tRNA), and other RNA species. Values represent the percentage of annotated reads assigned to each biotype.

**(B)** Representative electropherograms showing the RNA size distribution of total RNA isolated from untreated WaGa and MKL-1 cells and their corresponding EVs. Cellular RNA displays prominent 18S and 28S rRNA peaks, whereas EV-derived RNA lacks detectable ribosomal RNA peaks.

**(C)** RNA-sequencing characteristics of WaGa-derived EVs under the indicated conditions, including the percentage of aligned reads, reads assigned to protein-coding genes, and reads mapping to exonic or intronic regions; values for parental-cell RNA are shown for comparison.

**Figure S5: Characterization of the RNA cargo of untreated MKL-1-derived EVs.**

**(A)** Volcano plot showing differentially abundant transcripts in untreated MKL-1-derived EVs compared with the corresponding parental cells. Transcripts were considered significantly enriched or depleted with a log_2_ fold change > 1 or < –1 and an adjusted *p* value (padj.) < 0.05. **(B)** Gene Ontology (GO) Biological Process enrichment analysis of significantly enriched and depleted transcripts identified in panel B was performed using DAVID. Among all significantly enriched GO terms (FDR ≤ 0.05), the 20 terms with the highest gene counts were selected and grouped according to their biological function. Circle size represents the number of associated genes. Red circles indicate GO terms enriched among transcripts with higher relative abundance in EVs, whereas blue circles indicate GO terms enriched among transcripts. **(C)** Motif enrichment analysis of transcripts enriched in untreated MKL-1-derived EVs. Enriched sequence motifs were identified using AME (FDR < 0.1). The corresponding RNA-binding proteins (RBPs), enriched motifs, and FDR values are shown.

**Figure S6: Characterization of the miRNA cargo of untreated MKL-1-derived EVs.**

**(A)** Distribution of small RNA sequencing reads across annotated small RNA biotypes in untreated MKL-1-derived EVs. Numbers indicate the total number of aligned reads and the relative contribution of each small RNA class to the annotated reads (%). **(B)** Manhattan plot showing normalized miRNA abundance (log_10_ reads per million, RPM) in untreated MKL-1 parental cells (left) and their corresponding EVs (right). Red dots indicate the 15 most abundant miRNAs in each sample type. Each dot represents one miRNA (n = 2). **(C)** Gene Ontology (GO) Biological Process enrichment analysis of experimentally validated target genes (miRTarBase) of the 15 most abundant EV-associated miRNAs was performed using DAVID. Among all significantly enriched GO terms (FDR ≤ 0.05), those with the highest gene counts were selected and grouped according to their biological function. Circle size represents the number of associated genes, and color intensity indicates the significance of enrichment (FDR ≤ 0.05).

**Figure S7: Network analysis identifies genes targeted by multiple EV-associated miRNAs.**

**(A)** Network analysis of experimentally validated target genes (miRTarBase) of the 15 most abundant EV-associated miRNAs. miRNAs are shown as white nodes, whereas target genes are color-coded according to the number of miRNAs targeting each gene. Edges indicate experimentally validated miRNA–target gene interactions. **(B)** Genes targeted by multiple miRNAs identified in the network analysis shown in panel A, together with their annotated biological functions. **(C)** Gene Ontology (GO) Biological Process enrichment analysis of the multi-targeted genes was performed using DAVID. Among all significantly enriched GO terms (FDR ≤ 0.05), those with the highest gene counts were selected and grouped according to their biological function. Circle size represents the number of associated genes, and color intensity indicates the significance of enrichment (FDR ≤ 0.05).

**Figure S8: Knockdown of the viral small T antigen induces limited changes in the EV-associated miRNA cargo. (A)** Volcano plot showing differentially abundant miRNAs in WaGa-derived EVs following sT knockdown (shRNA sT Dox) compared with the corresponding DMSO-treated control (shRNA sT DMSO). miRNAs with a log_2_ fold change > 1 or < –1 and an adjusted *p* value (padj.) < 0.05 were considered significantly enriched or depleted. Significantly enriched miRNAs are shown in red, significantly depleted miRNAs in blue, and non-significant miRNAs in grey. **(B)** Hierarchical clustering heatmap of normalized miRNA abundances in EVs derived from untreated WaGa cells (wt), shRNA sT cells treated with DMSO or doxycycline (Dox), and shRNA scramble (scr) cells treated with DMSO or doxycycline. Rows represent individual miRNAs and columns represent biological replicates. Colors indicate row Z-scores of normalized miRNA abundance. **(C)** Gene Ontology (GO) Biological Process enrichment analysis of experimentally validated target genes (miRTarBase) of the significantly altered miRNAs identified in panel A was performed using DAVID. Among all significantly enriched GO terms (FDR ≤ 0.05), those with the highest gene counts were selected and grouped according to their biological function. Circle size represents the number of associated genes, and color intensity indicates the significance of enrichment (−log10(FDR)).

**Figure S9: Gene Ontology enrichment analysis of WaGa- and MKL-1-parental cells.**

**(A-B)** Gene Ontology (GO) Biological Process enrichment analysis of transcripts identified in untreated WaGa-derived EVs (A) and MKL-1-derived EVs (B) using DAVID. Among all significantly enriched GO terms (FDR ≤ 0.05), the terms with the highest gene counts were selected and grouped according to their biological function. Circle size represents the number of associated genes, and color intensity indicates the significance of enrichment (−log10(FDR)).

**Figure S10: Knockdown of the viral small T antigen induces limited changes in the EV RNA cargo.**

**(A)** Hierarchical clustering heatmap of transcript abundances in WaGa-derived EVs isolated from untreated cells (wt), shRNA sT cells treated with DMSO or doxycycline (Dox), and shRNA scramble (scr) cells treated with DMSO or doxycycline. Rows represent transcripts and columns represent individual biological replicates. Colors indicate row Z-scores of normalized transcript abundance. **(B)** Hierarchical clustering heatmap of transcript abundances in EVs derived from MKL-1 cells under the corresponding experimental conditions. Rows represent transcripts and columns represent individual biological replicates. Colors indicate row Z-scores of normalized transcript abundance. **(B)** Heatmap showing the relative abundance of transcripts identified as differentially abundant in extracellular vesicles (EVs) following small T antigen (sT) knockdown. In contrast to the heatmap shown in (A), which includes parental-cell samples, this analysis is restricted to EV samples to facilitate comparison of the effects of sT knockdown on EV-associated transcripts. **(C)** Volcano plot showing differentially abundant transcripts in WaGa-derived EVs following sT knockdown (shRNA sT Dox, sT OFF) compared with the corresponding DMSO-treated control (shRNA sT DMSO, sT ON). Transcripts with a log2fold change > 1 or < –1 and an adjusted *p* value (padj.) < 0.05 were considered significantly enriched or depleted. Fifteen transcripts showed significantly higher relative abundance in EVs following sT knockdown, whereas no significantly depleted transcripts were identified.

**Figure S11: Knockdown of the viral small T antigen induces only minor changes in the proteome of MKL-1-derived EVs. (A)** Hierarchical clustering heatmap of protein abundances in EVs isolated from MKL-1 cells carrying shRNA sT or shRNA scramble (scr) following treatment with DMSO or doxycycline (Dox). Rows represent individual proteins and columns represent biological replicates. Colors indicate relative protein abundance (row Z-score). **(B)** Volcano plot showing differentially abundant proteins in EVs derived from sT knockdown cells (shRNA sT Dox, sT OFF) compared with the corresponding DMSO-treated control (shRNA sT DMSO, sT ON). Proteins with a log2fold change > 1 or < –1 and an adjusted *p* value (padj.) < 0.05 were considered significantly enriched or depleted. Two proteins (KRT5 and KRT14) exhibited significantly higher relative abundance following sT knockdown, whereas no proteins showed significantly lower relative abundance.

## Notes

### Competing Interest Statement

The authors have declared no competing interest.

