## Supplementary figures and images for "Multi-omics characterization of extracellular vesicles derived from virus-positive Merkel cell carcinoma cells"

### Supplementary Figure 1

#### Slide 1
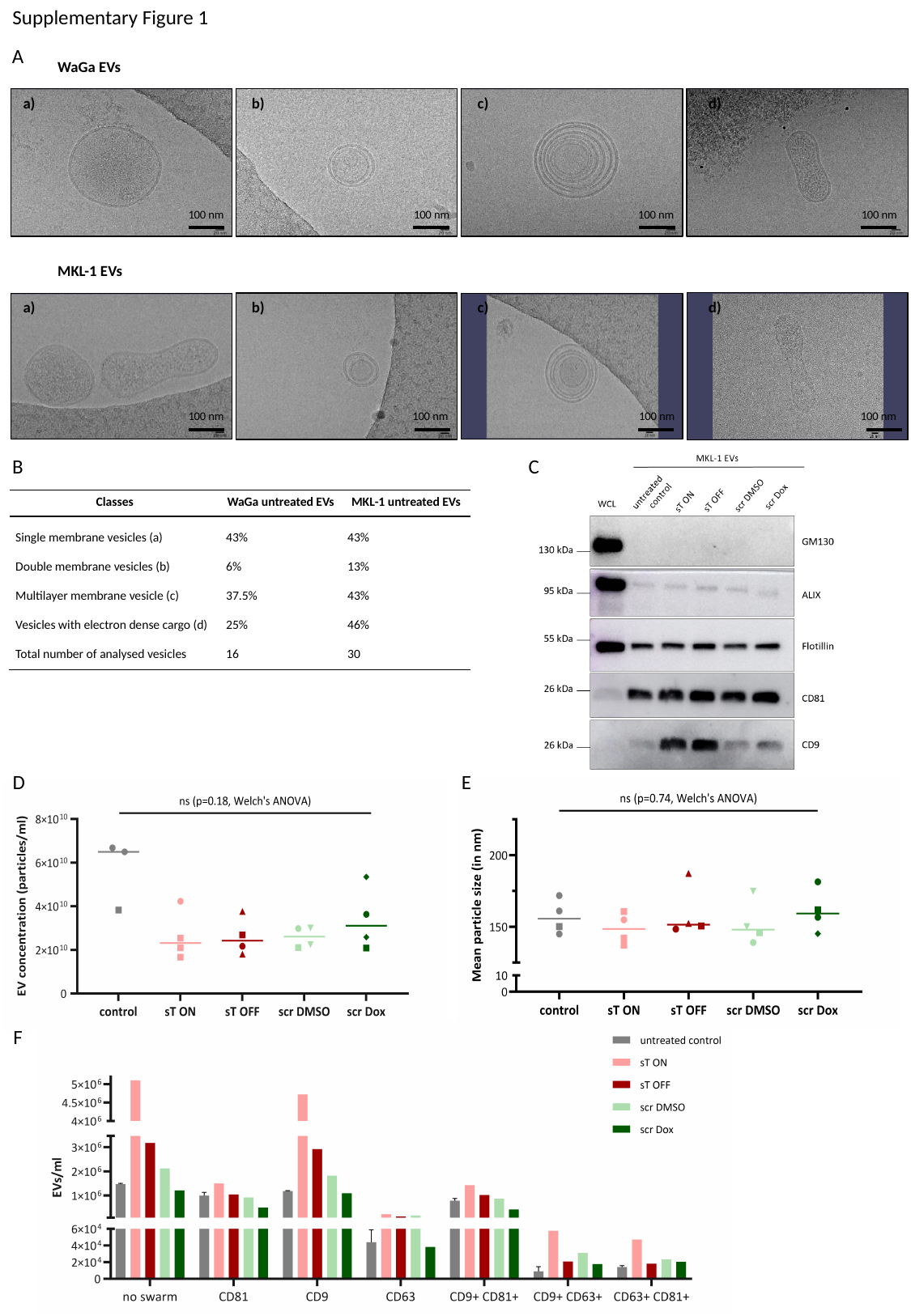

Supplementary Figure 1
A
 WaGa EVs
a) b) c) d)
100 nm
100 nm
100 nm
100 nm
 MKL-1 EVs
a) b) c) d)
100 nm
100 nm
100 nm
100 nm
B
C
D
E
F

### Supplementary Figure 2

#### Slide 1
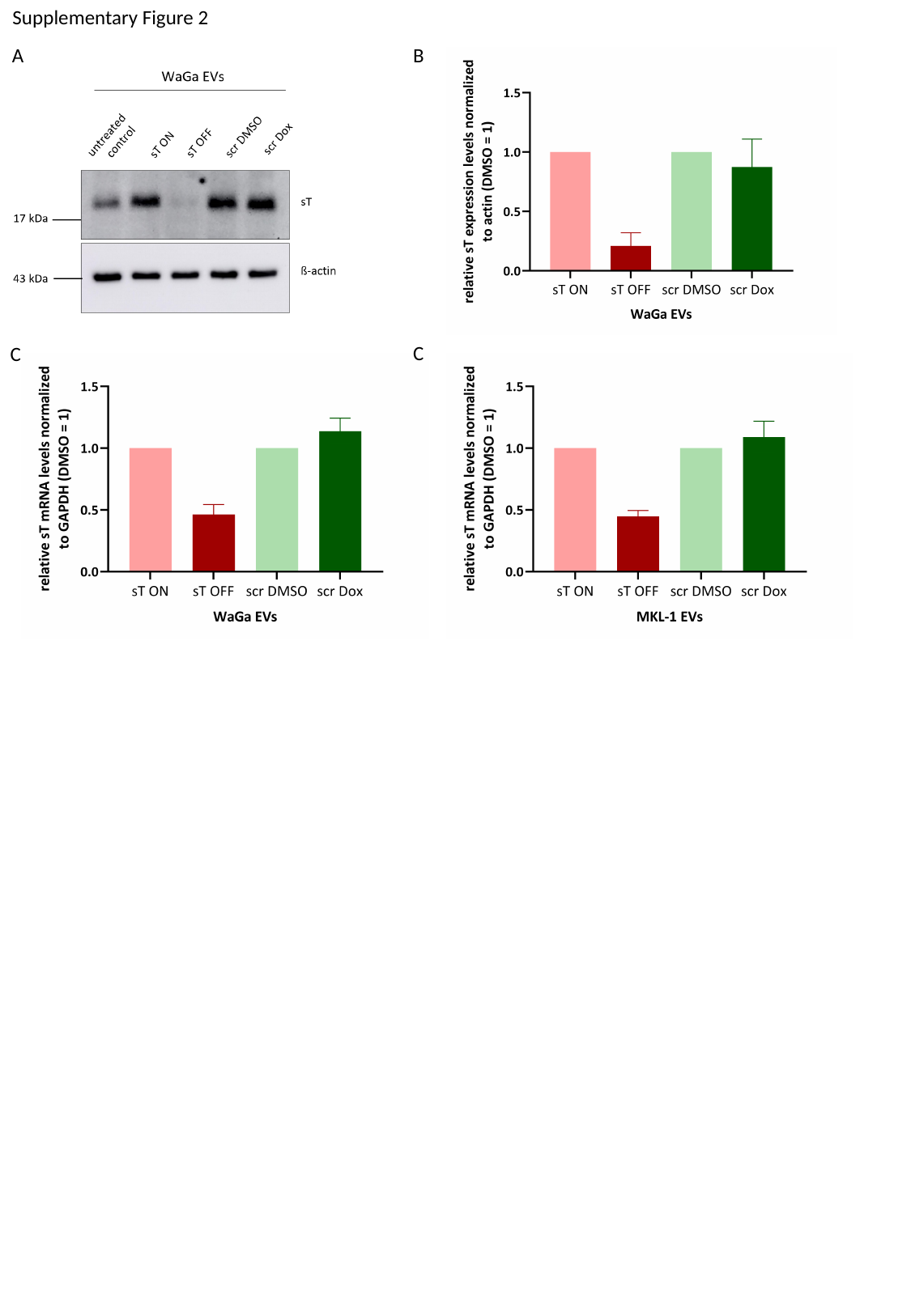

Supplementary Figure 2
B
A
C
C

### Supplementary Figure 3

#### Slide 1
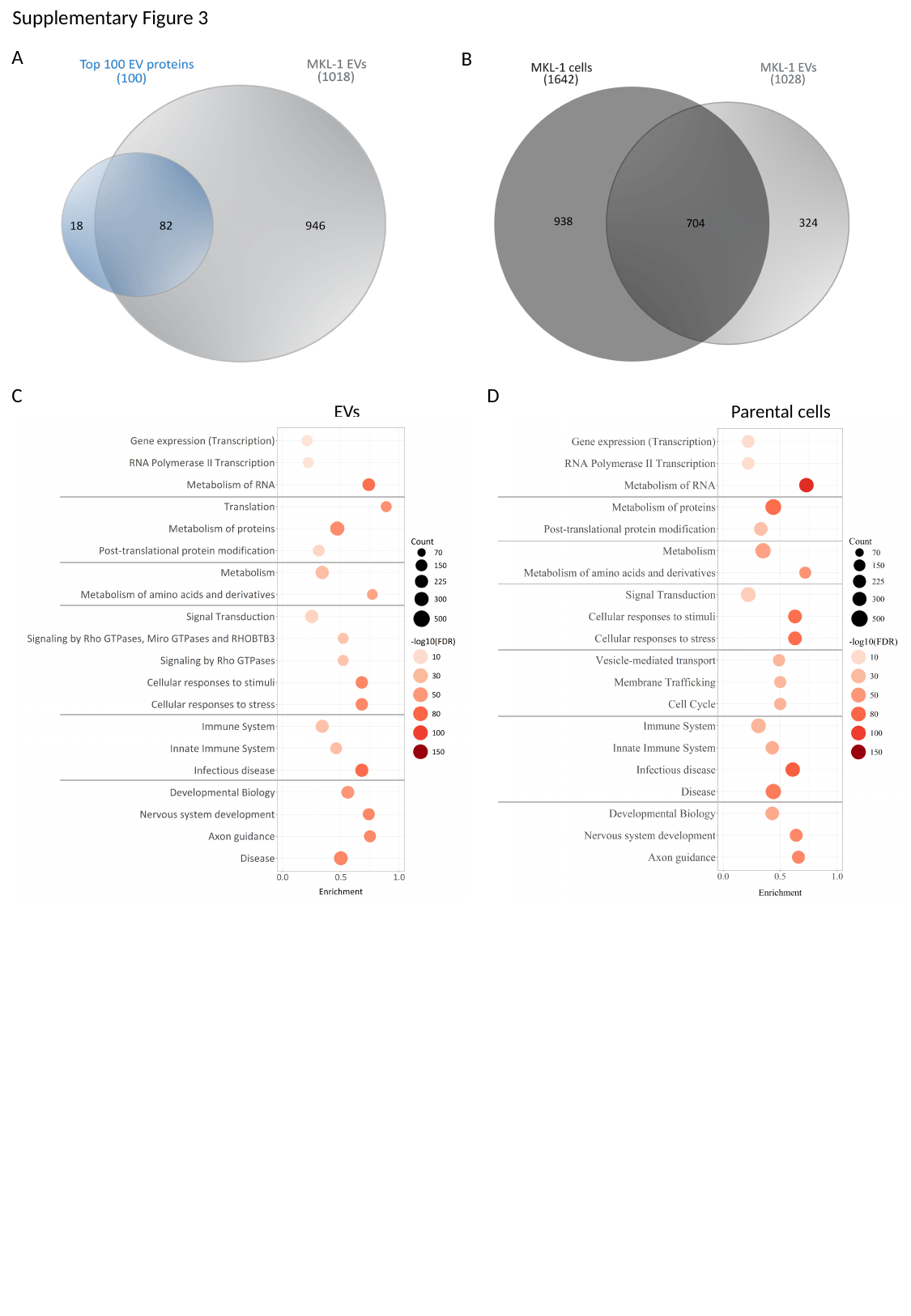

Supplementary Figure 3
A
B
C
D
EVs
Parental cells

### Supplementary Figure 6

#### Slide 1
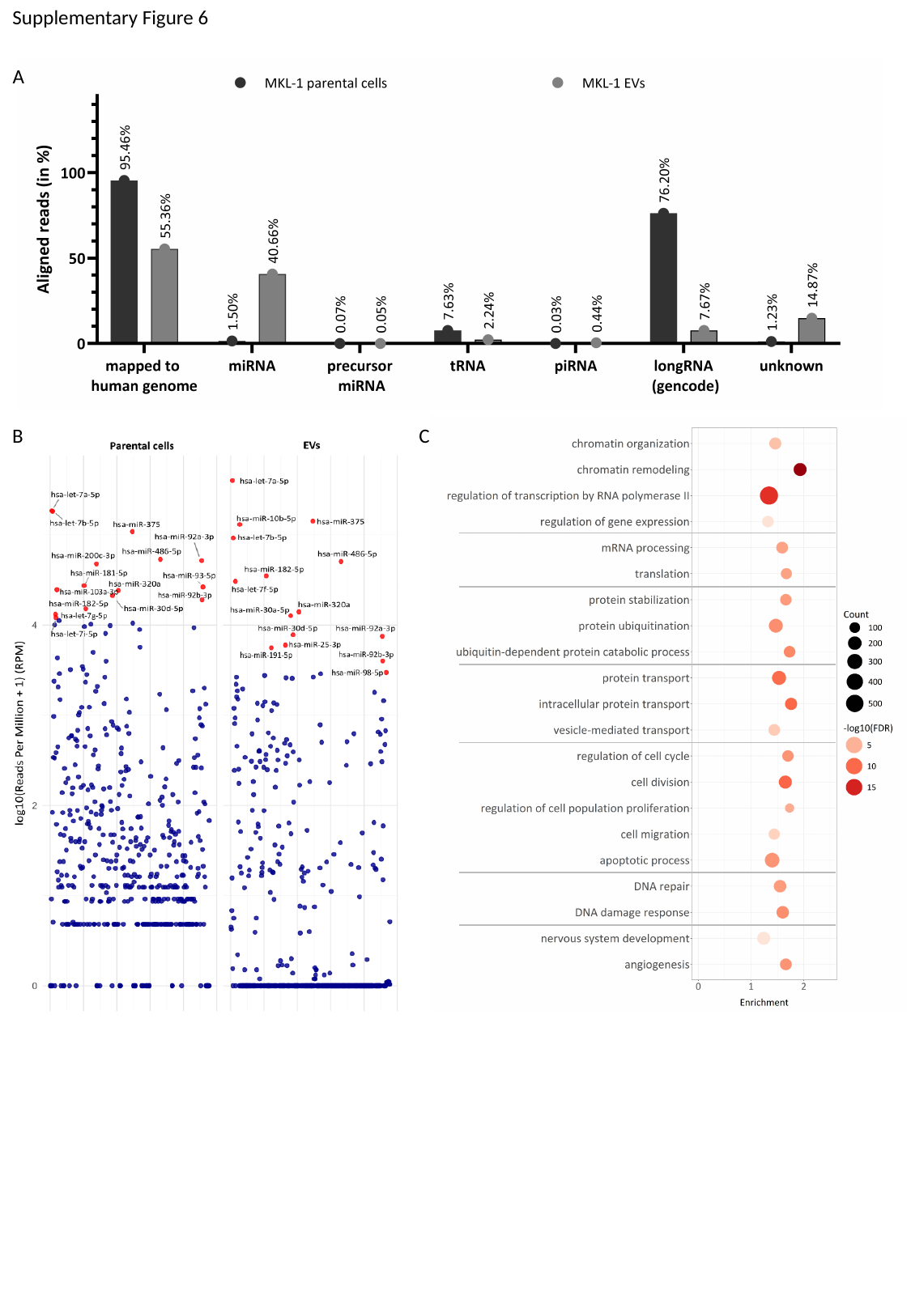

Supplementary Figure 6
A
C
B

### Supplementary Figure 9

#### Slide 1
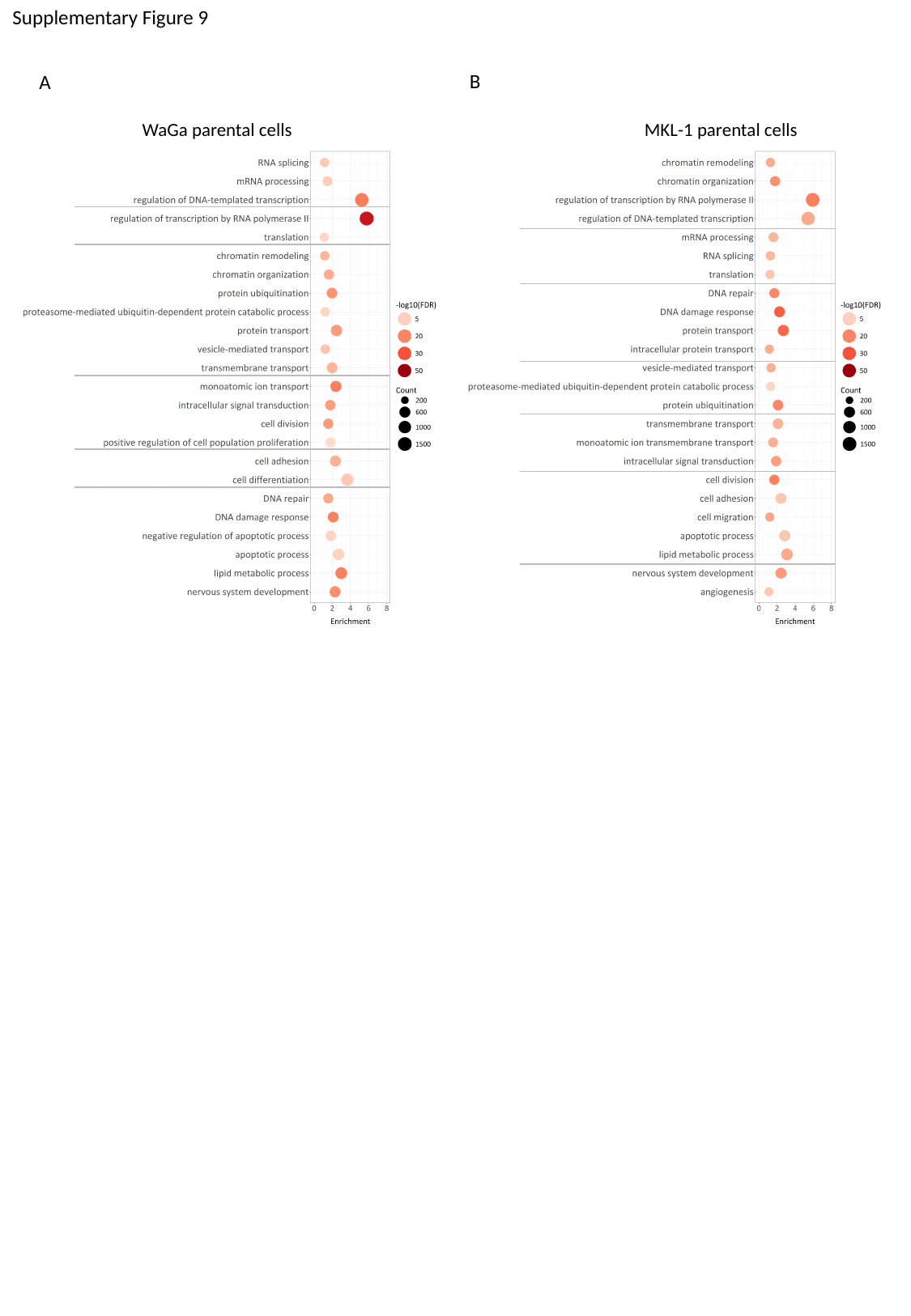

Supplementary Figure 9
B
A
WaGa parental cells
MKL-1 parental cells

### Supplementary Figure 10

#### Slide 1
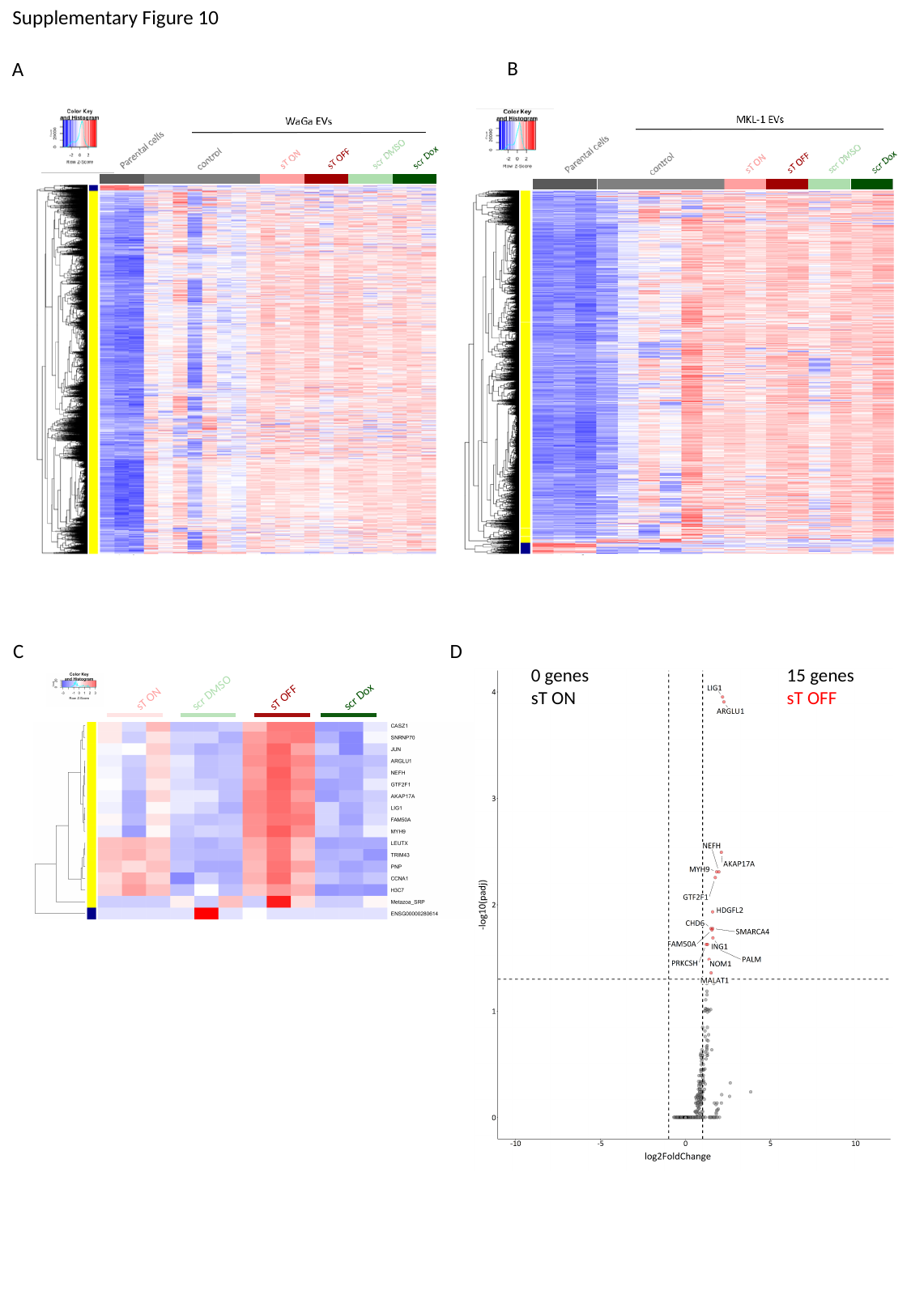

Supplementary Figure 10
B
A
C
D
15 genes
sT OFF
0 genes
sT ON
scr DMSO
sT ON
sT OFF
scr Dox
