## Supplementary Figure 4 for "Multi-omics characterization of extracellular vesicles derived from virus-positive Merkel cell carcinoma cells"

### Slide 1
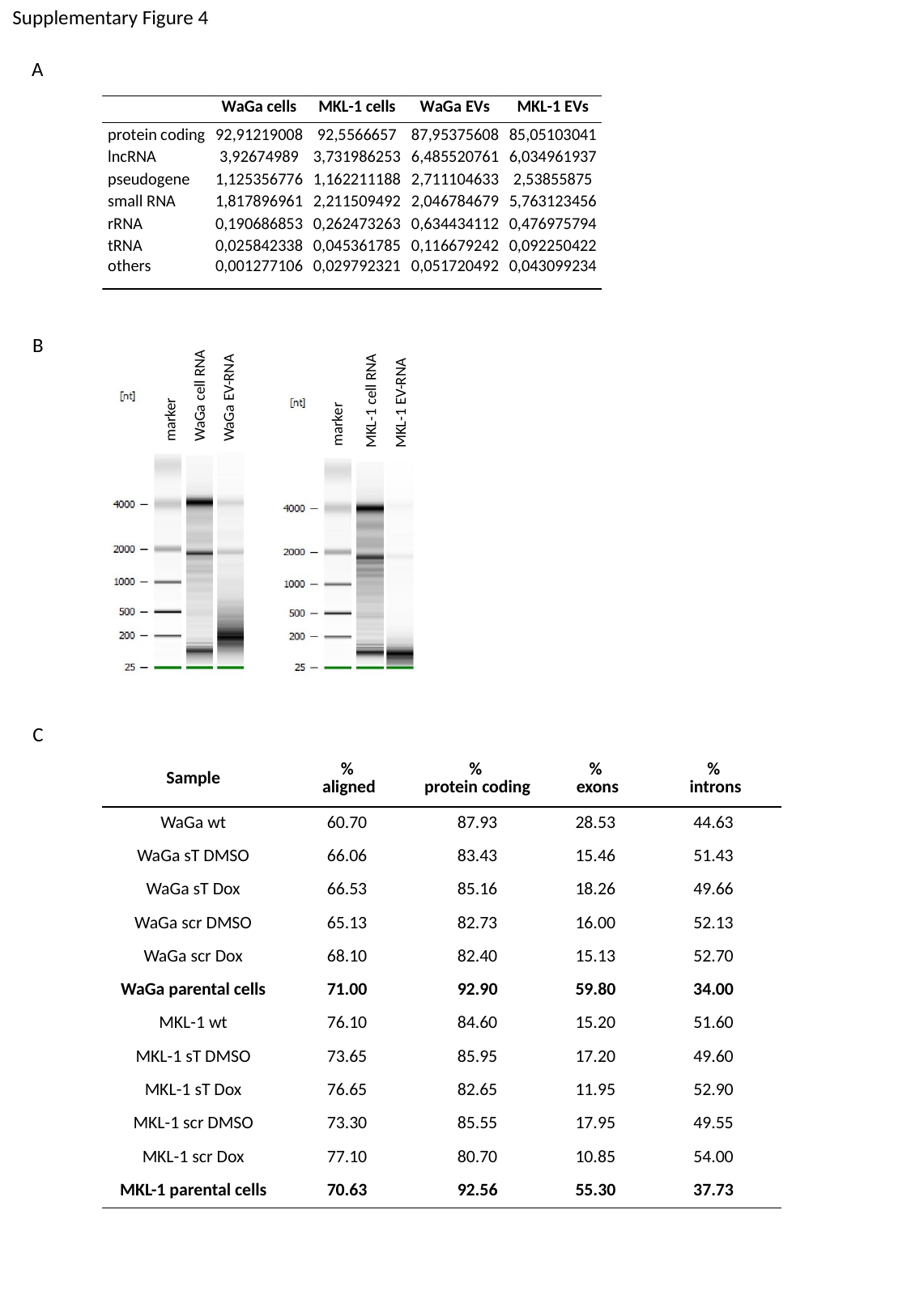

Supplementary Figure 4
A
WaGa cell RNA
WaGa EV-RNA
MKL-1 EV-RNA
MKL-1 cell RNA
B
marker
marker
C
| Sample | % aligned | % protein coding | % exons | % introns |
| --- | --- | --- | --- | --- |
| WaGa wt | 60.70 | 87.93 | 28.53 | 44.63 |
| WaGa sT DMSO | 66.06 | 83.43 | 15.46 | 51.43 |
| WaGa sT Dox | 66.53 | 85.16 | 18.26 | 49.66 |
| WaGa scr DMSO | 65.13 | 82.73 | 16.00 | 52.13 |
| WaGa scr Dox | 68.10 | 82.40 | 15.13 | 52.70 |
| WaGa parental cells | 71.00 | 92.90 | 59.80 | 34.00 |
| MKL-1 wt | 76.10 | 84.60 | 15.20 | 51.60 |
| MKL-1 sT DMSO | 73.65 | 85.95 | 17.20 | 49.60 |
| MKL-1 sT Dox | 76.65 | 82.65 | 11.95 | 52.90 |
| MKL-1 scr DMSO | 73.30 | 85.55 | 17.95 | 49.55 |
| MKL-1 scr Dox | 77.10 | 80.70 | 10.85 | 54.00 |
| MKL-1 parental cells | 70.63 | 92.56 | 55.30 | 37.73 |
