## Supplementary Figure 5 for "Multi-omics characterization of extracellular vesicles derived from virus-positive Merkel cell carcinoma cells"

### Slide 1
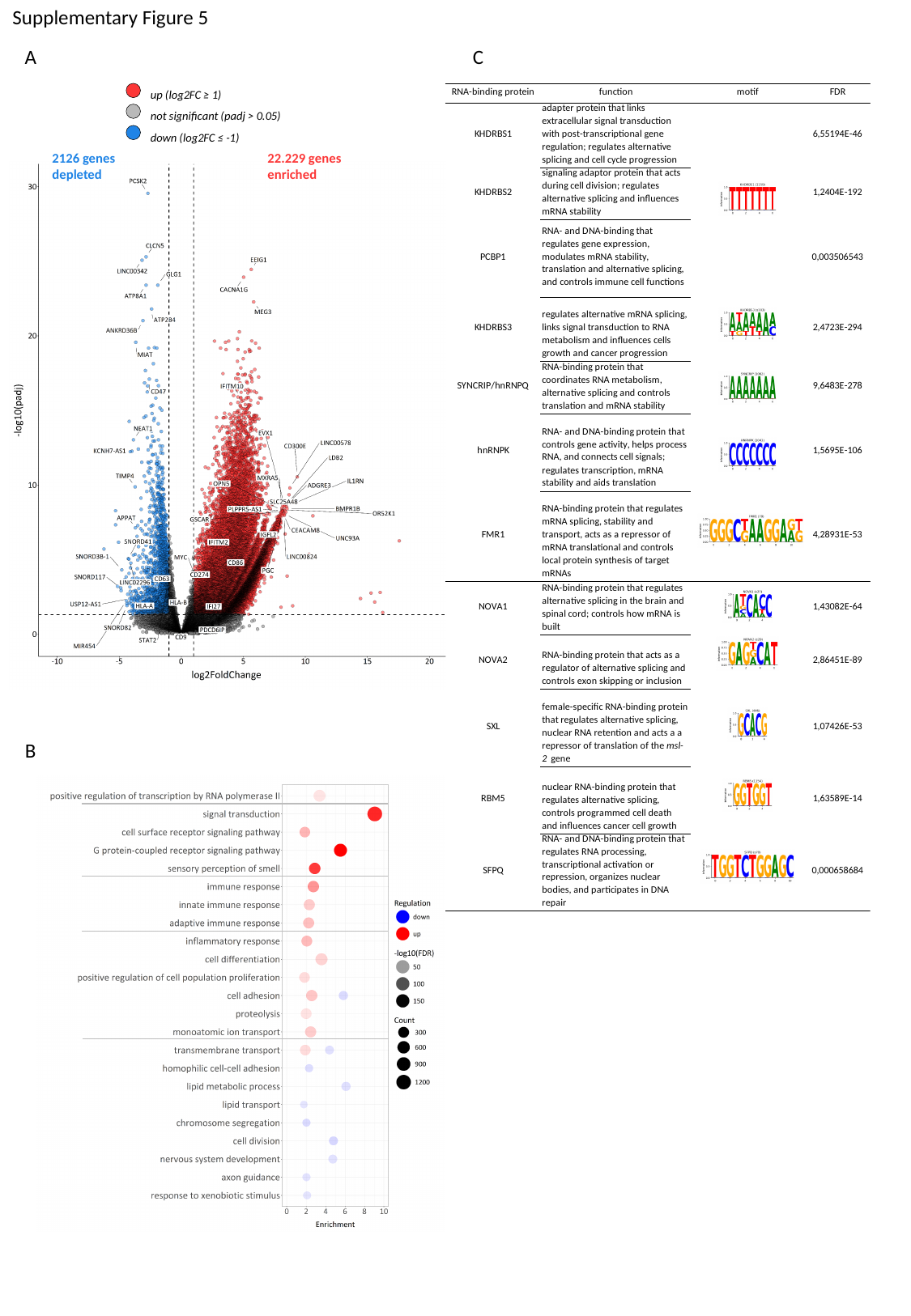

Supplementary Figure 5
C
A
up (log2FC ≥ 1)
not significant (padj > 0.05)
down (log2FC ≤ -1)
2126 genes
depleted
22.229 genes
enriched
B
