## Supplementary Figure 7 for "Multi-omics characterization of extracellular vesicles derived from virus-positive Merkel cell carcinoma cells"

### Slide 1
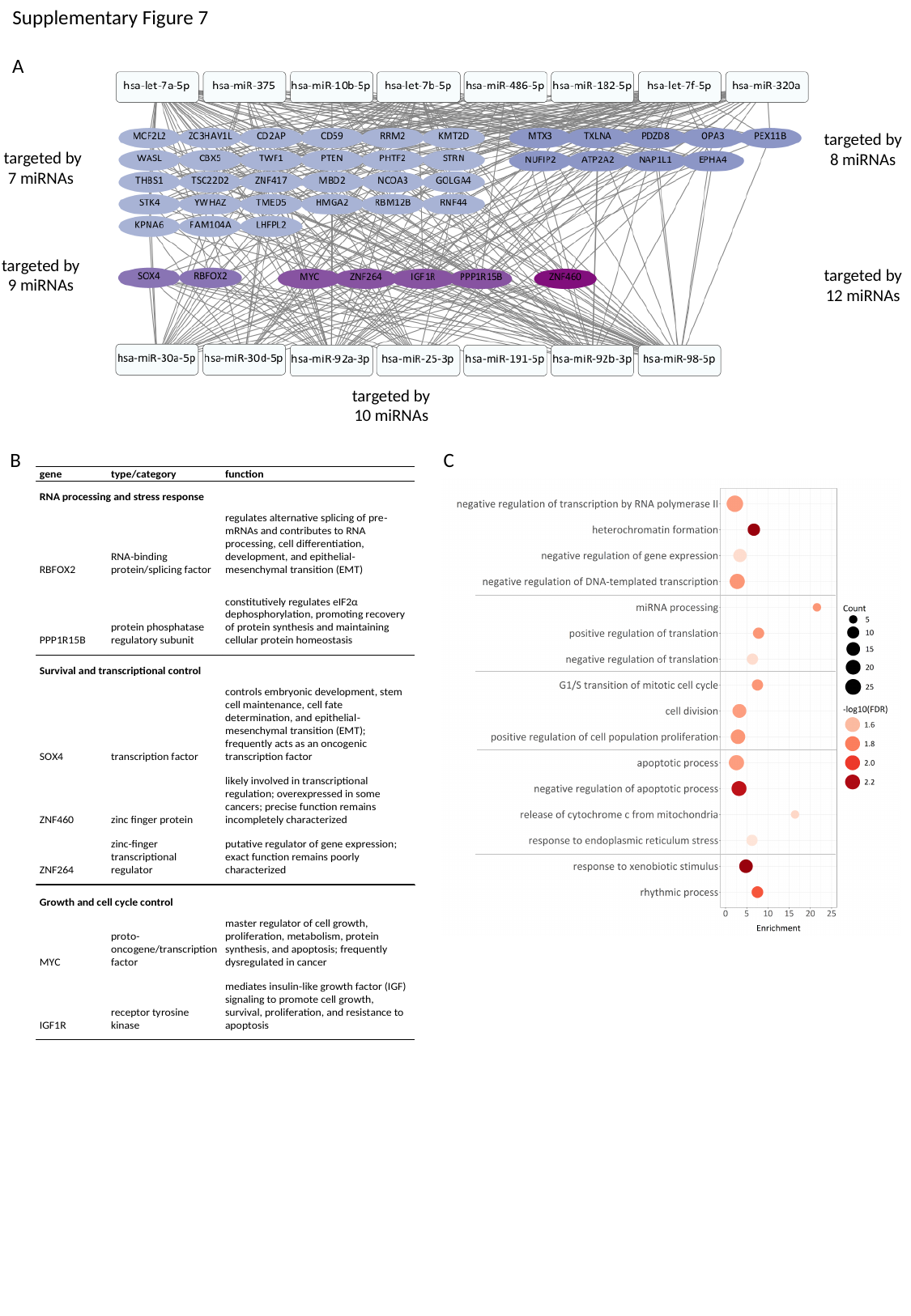

Supplementary Figure 7
A
targeted by
8 miRNAs
targeted by
7 miRNAs
targeted by
9 miRNAs
targeted by
12 miRNAs
targeted by
10 miRNAs
B
C
