## Supplementary Figure 8 for "Multi-omics characterization of extracellular vesicles derived from virus-positive Merkel cell carcinoma cells"

### Slide 1
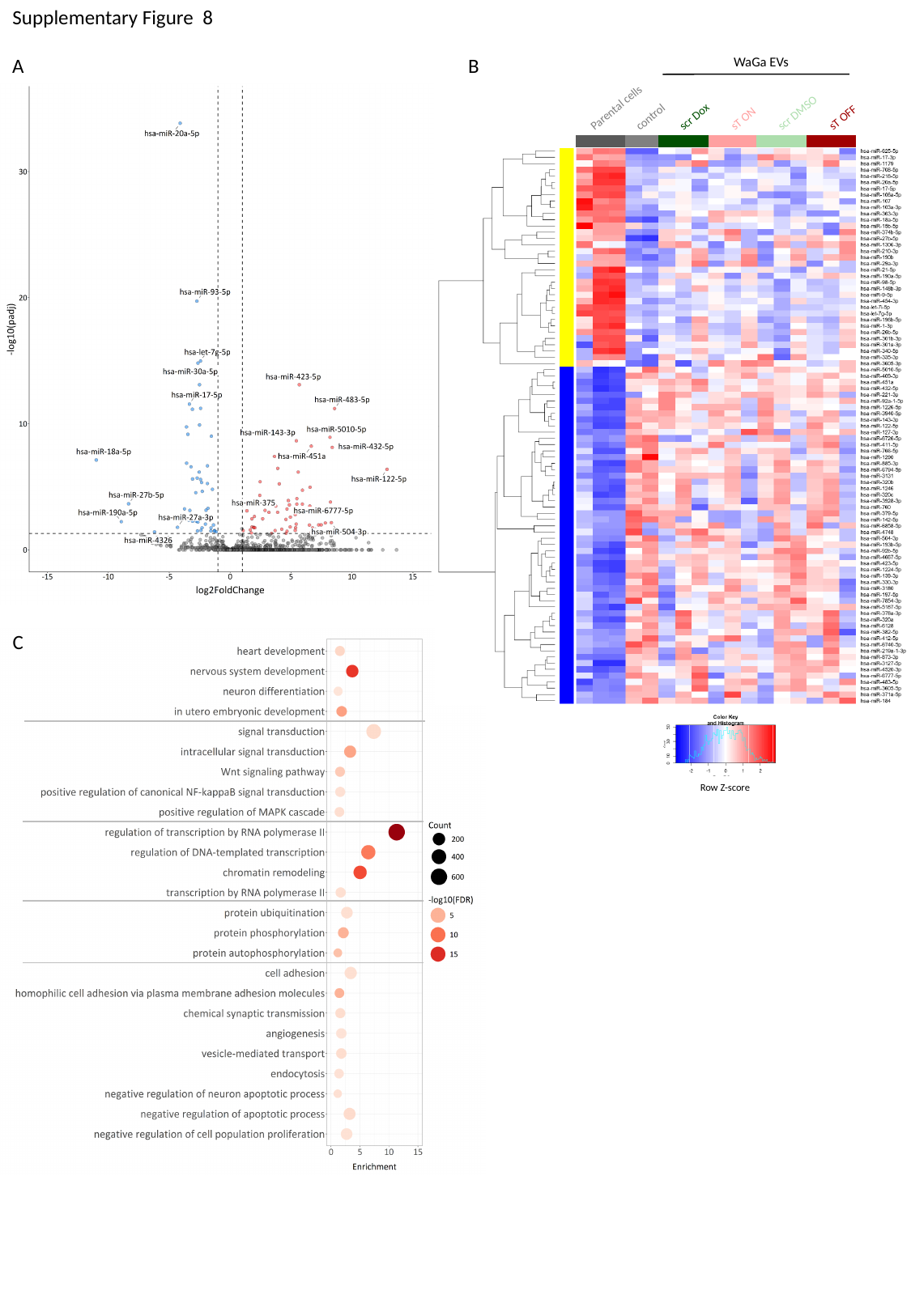

Supplementary Figure 8
WaGa EVs
Parental cells
scr DMSO
control
sT ON
sT OFF
scr Dox
Row Z-score
B
A
C
