## Supplementary Figure 11 for "Multi-omics characterization of extracellular vesicles derived from virus-positive Merkel cell carcinoma cells"

### Slide 1
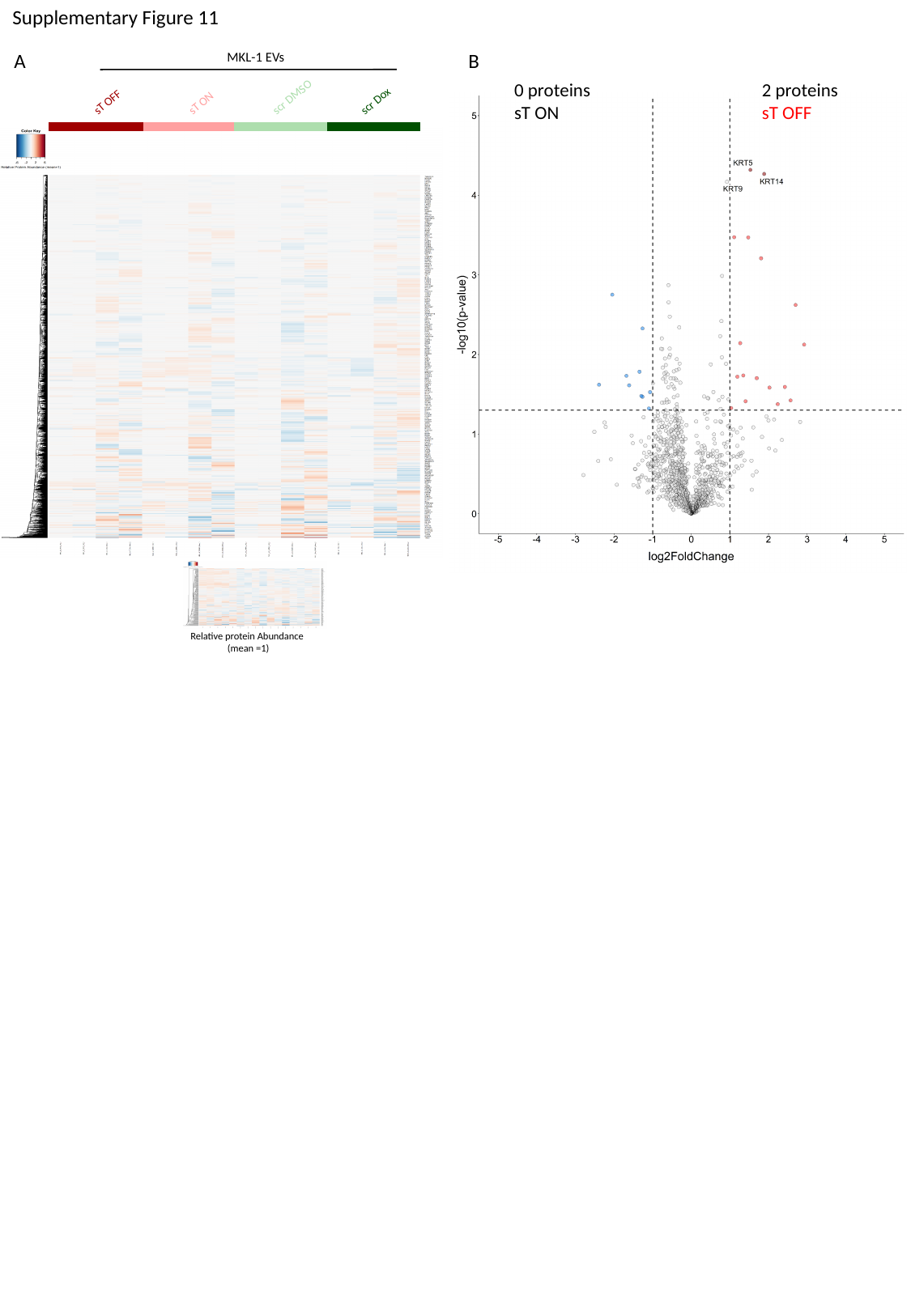

Supplementary Figure 11
MKL-1 EVs
scr DMSO
sT OFF
sT ON
scr Dox
A
B
0 proteins
sT ON
2 proteins
sT OFF
Relative protein Abundance
(mean =1)
