## Supplementary Table 1 for "Multi-omics characterization of extracellular vesicles derived from virus-positive Merkel cell carcinoma cells"

**Supplementary Table S1:**

**Top 100 most abundant proteins identified in extracellular vesicles released by WaGa and MKL-1 cells,** columns are sorted in alphabetical order

| **Top 100 proteins identified in WaGa EVs** | **Top 100 proteins identified in MKL-1 EVs** |
| --- | --- |
| A2M | A2M |
| ACLY | ACLY |
| ACTB | ACTB |
| ACTN1 | ACTN1 |
| ACTN4 | ACTN4 |
| ADAM10 | ADAM10 |
| AHCY | AHCY |
| ALB | ALB |
| ALDOA | ALDOA |
| ANXA2 | ANXA2 |
| ANXA5 | ANXA5 |
| ANXA6 | ANXA6 |
| ANXA7 | ANXA7 |
| ATP1A1 | ATP1A1 |
| CCT2 | CAP1 |
| CCT3 | CCT2 |
| CCT4 | CCT3 |
| CCT6A | CCT4 |
| CCT8 | CCT6A |
| CD81 | CCT8 |
| CD9 | CD9 |
| CFL1 | CFL1 |
| CLTC | CLIC1 |
| EEF1A1 | CLTC |
| EEF1G | EEF1A1 |
| EEF2 | EEF1G |
| EIF4A1 | EEF2 |
| ENO1 | EIF4A1 |
| EZR | ENO1 |
| FASN | EZR |
| FLNA | FASN |
| FN1 | FLNA |
| GAPDH | FLOT1 |
| GDI2 | FLOT2 |
| GNAS | FN1 |
| GNB1 | GAPDH |
| GNB2 | GDI2 |
| GPI | GNAI2 |
| HIST1H4A | GNAS |
| HSP90AA1 | GNB1 |
| HSP90AB1 | GNB2 |
| HSPA5 | GPI |
| HSPA8 | GSN |
| KPNB1 | HIST1H4A |
| LDHA | HSP90AA1 |
| LDHB | HSP90AB1 |
| LGALS3BP | HSPA5 |
| MFGE8 | HSPA8 |
| MYH9 | IQGAP1 |
| PFN1 | ITGB1 |
| PGAM1 | KPNB1 |
| PGK1 | LDHA |
| PKM | LDHB |
| PPIA | LGALS3BP |
| PRDX1 | MFGE8 |
| PRDX2 | MSN |
| RAB10 | MYH9 |
| RAB5C | PDCD6IP |
| RAB7A | PFN1 |
| RAP1B | PGAM1 |
| RHOA | PGK1 |
| SLC3A2 | PKM |
| TFRC | PPIA |
| TLN1 | PRDX1 |
| TPI1 | PRDX2 |
| TUBB4B | RAB10 |
| VCP | RAB7A |
| YWHAB | RAP1B |
| YWHAE | RHOA |
| YWHAG | SLC3A2 |
| YWHAQ | TCP1 |
| YWHAZ | TFRC |
|  | TLN1 |
|  | TPI1 |
|  | TUBB4B |
|  | VCL |
|  | VCP |
|  | YWHAB |
|  | YWHAE |
|  | YWHAG |
|  | YWHAQ |
|  | YWHAZ |
