## Supplementary Table 2 for "Multi-omics characterization of extracellular vesicles derived from virus-positive Merkel cell carcinoma cells"

**Supplementary Table S2:**

Reactome pathway enrichment analysis of the WaGa-derived extracellular vesicle proteome shown in Figure 2C. The 20 most significantly enriched pathways are shown.

| **Reactome Pathway** | **Reactome term ID** | **Gene Count** | **Adjusted p-value (FDR)** | **proteins identified in this pathway** |
| --- | --- | --- | --- | --- |
| Gene expression (Transcription) | HSA-74160 | 75 | 0,00017 | PSMC4, CBX5, SNRPD3, SUPT16H, CSNK2A1, CAT, SRSF6, H4C6, YWHAH, APOE, TRIM28, RPA1, H3-3B, SRSF1, THBS1, PSMA6, PRDX1, YWHAE, PRDX5, SNRPF, PSMB6, PSMA5, RPS27A, SSRP1, SMARCA5, CALM3, U2AF1, NPM1, PRDX2, H2AC8, YWHAG, H2AZ2, PSMD2, PSMC5, PRMT5, PPP2R1A, SRSF7, SRRM1, H2AC20, HSP90AA1, CBX3, HSPD1, SFN, BAZ1B, PSMB5, PARP1, PSMD4, H2BC21, SRSF11, TP53RK, YBX1, YWHAB, RBBP4, SRSF3, SRSF4, CSNK2B, PCNA, YWHAQ, SRSF2, YWHAZ, SNRPG, SNRPE, GPI, SNRPB, DDX39B, TFAM, HDAC2, RAN, H2BC14, SRRT, PSMC3, PSMC6, ACTB, CTNNB1, EIF4A3, DEK |
| Metabolism of RNA | HSA-8953854 | 147 | 7,62E-69 | PSMC4, SNRPD3, RTCB, PNN, HNRNPL, FBL, RPL18A, RPL19, DDX5, RPS12, SRSF6, HSPB1, RBM39, SNRPA1, SRSF1, RPL35, PPIG, PSMA6, RBM25, RPL8, ACIN1, CCAR1, SNRPF, PSMB6, RPS11, PSMA5, RPS27A, RPS3, U2AF1, EIF4A1, RPL29, RPS23, SRRM2, PCBP1, RPL13, RPL15, RPL38, PSMD2, PSMC5, RPL4, PABPC1, HNRNPD, SNRNP200, NCL, RPS15A, SF3B2, PRMT5, PPP2R1A, HNRNPM, SRSF7, SRRM1, TNPO1, DHX15, RPL7, HNRNPA1, RPS2, SNRPD2, RPL3, RPL36AL, RPL27A, RPL21, RPS3A, RPL22, HNRNPA2B1, RPS26, HNRNPH1, PTBP1, PSMB5, DHX9, PSMD4, BCAS2, DKC1, RPL5, RBMX, SRSF11, RPL7A, TP53RK, YBX1, SET, YWHAB, RPS4X, SRSF3, SRSF4, RPL10A, HNRNPR, HSPA1B, HNRNPK, RPS16, RPS6, NOP56, RPS9, RPL13A, SRSF2, HNRNPA3, RPL24, RPL34, YWHAZ, RPL14, RPS8, KHSRP, RPS14, RPL23A, RPSA, EFTUD2, RPS18, SNRPG, SNRPE, RPL6, SNRPB, RPL10, RPS24, DDX39B, RPL32, TRA2B, ANP32A, RPL37A, SRSF10, RPL23, RPS20, RPLP2, RPS25, RPS13, HSPA8, RAN, RPL18, HNRNPC, RPL28, RPL26, RPL17, RPL36, RPL27, RPS15, LSM4, RPS19, RPS28, RPS5, SNRNP70, SRRT, PSMC3, RPS10, PSMC6, HNRNPU, RPS7, RPL35A, RPL11, EIF4A3, RPS27 |
| Translation | HSA-72766 | 85 | 1,01E-48 | EIF3D, RPL18A, RPL19, RPS12, EIF2S1, NARS1, RPL35, RPL8, DARS1, RPS11, RPS27A, RPS3, EIF4A1, RPL29, RPS23, RPL13, EEF2, RPL15, RPL38, RPL4, PABPC1, RPS15A, EEF1G, EIF3C, EEF1A1, RPL7, RPS2, RPL3, RPL36AL, RPL27A, RPL21, RPS3A, RPL22, RPS26, EIF3B, RPL5, RPL7A, PPA1, YARS1, EIF3I, RPS4X, RPL10A, IARS1, VARS1, RPS16, EEF1E1, RPS6, RPS9, RPL13A, RPL24, LARS1, RPL34, AIMP1, RPL14, RPS8, RPS14, RPL23A, RPSA, RPS18, RPL6, EEF1D, RPL10, RPS24, RPL32, RPL37A, RPL23, RPS20, RPLP2, RPS25, RPS13, RPL18, RPL28, RPL26, RPL17, RPL36, RPL27, RPS15, RPS19, RPS28, RPS5, RPS10, RPS7, RPL35A, RPL11, RPS27 |
| Metabolism of proteins | HSA-392499 | 192 | 7,06E-47 | PSMC4, CBX5, NANS, EIF3D, CSNK2A1, TBCB, RPL18A, RPL19, DDX5, RAB35, RPS12, LTF, H4C6, CANX, TUBA4A, LMAN1, APOE, TRIM28, RPA1, H3-3B, EIF2S1, ARF3, NARS1, CCT7, RPL35, TUBB2B, THBS1, ADAM10, RAB21, LYZ, UBE2K, PSMA6, PFDN1, RAB11A, RPL8, CAPZA1, DARS1, RAB10, RAB7A, MFGE8, PSMB6, RPS11, PSMA5, RPS27A, AHSG, CCT6A, RPS3, CCT5, UCHL1, XPC, CCT8, EIF4A1, RPL29, CCT3, ALB, RPS23, NPM1, CCT2, HSP90B1, PDIA3, DDB1, H2AC8, GNB2, KIF5B, RPL13, EEF2, RPL15, RPL38, PSMD2, PSMC5, RPL4, RAB6A, PABPC1, PRKDC, RUVBL1, RPS15A, TUBB3, CALR, SEC16A, P4HB, EEF1G, H2AC20, EIF3C, H2AC21, UBA1, EEF1A1, RPL7, PARK7, GANAB, TUBB4B, RPS2, RPL3, RPL36AL, RPL27A, RPL21, RPS3A, RPL22, FN1, RPS26, DYNC1H1, VCP, EIF3B, TOP1, PSMB5, PARP1, PFDN2, PSMD4, H2BC21, RPL5, RPL7A, TSPAN14, VDAC2, PPA1, YARS1, EIF3I, RPS4X, RAB14, RPL10A, IARS1, VARS1, CSNK2B, HNRNPK, RPS16, GNB1, PCNA, EEF1E1, TUBB2A, RPS6, NOP56, RPS9, RPL13A, RPL24, LARS1, CCT4, RPL34, AIMP1, RPL14, RPS8, DDX17, NSF, PDIA6, RPS14, RAB1A, RPL23A, RPSA, RPS18, RHOA, CAPZB, RPL6, SUMO2, EEF1D, TOP2A, RPL10, RPS24, RPL32, RPL37A, VAMP2, RPL23, RPS20, HDAC2, RPLP2, RPS25, RPS13, HSPA8, CALU, TUBA1A, RPL18, RAB5C, HNRNPC, RPL28, RPL26, RPL17, RPL36, RPL27, RPS15, PRKCSH, RPS19, RPS28, RPS5, H2BC14, PSMC3, RPS10, PSMC6, SPTAN1, ACTB, RPS7, CTNNB1, RPL35A, RPL11, PCMT1, CD59, RPS27 |
| Post-translational protein modification | HSA-597592 | 88 | 1,02E-08 | PSMC4, CBX5, NANS, DDX5, RAB35, H4C6, CANX, TUBA4A, LMAN1, APOE, TRIM28, RPA1, ARF3, TUBB2B, THBS1, ADAM10, RAB21, UBE2K, PSMA6, RAB11A, CAPZA1, RAB10, RAB7A, MFGE8, PSMB6, PSMA5, RPS27A, AHSG, UCHL1, XPC, ALB, NPM1, HSP90B1, PDIA3, DDB1, H2AC8, EEF2, PSMD2, PSMC5, RAB6A, PRKDC, RUVBL1, TUBB3, CALR, SEC16A, P4HB, H2AC20, H2AC21, UBA1, EEF1A1, PARK7, GANAB, TUBB4B, RPS2, FN1, DYNC1H1, VCP, TOP1, PSMB5, PARP1, PSMD4, H2BC21, VDAC2, RAB14, HNRNPK, PCNA, TUBB2A, DDX17, NSF, PDIA6, RAB1A, RHOA, CAPZB, SUMO2, TOP2A, HDAC2, HSPA8, CALU, TUBA1A, RAB5C, HNRNPC, PRKCSH, H2BC14, PSMC3, PSMC6, SPTAN1, ACTB, CD59 |
| Metabolism | HSA-1430728 | 164 | 4,20E-27 | PSMC4, UQCRC1, CSNK2A1, AHCY, RPL18A, RPL19, TPI1, RPS12, UMPS, APOE, SNAP25, ARF3, RPL35, PSMA6, SLC7A5, ALDH2, ATP5F1B, RPL8, GPC1, DARS1, AGPS, ESYT1, PSMB6, RPS11, PSMA5, RPS27A, GLUD1, RPS3, KPNB1, RPL29, ALB, RPS23, FABP5, INPPL1, CKB, FASN, DTYMK, GNB2, RPL13, PRSS1, RPL15, RPL38, PSMD2, PSMC5, RPL4, RPS15A, PKM, TALDO1, PPP2R1A, MDH2, IDH2, BSG, HSP90AA1, RPL7, RPS2, SERINC1, VAPA, RPL3, RPL36AL, RPL27A, RPL21, RPS3A, RPL22, THRAP3, ALDH9A1, DHCR7, RPS26, PSMB5, PSMD4, RPL5, PGAM1, GNAS, HSP90AB1, RPL7A, SDC4, PPA1, STXBP1, PGK1, GLO1, RPS4X, RAB14, RPL10A, SLC44A1, IARS1, CSNK2B, SLC3A2, APRT, RPS16, GNB1, ESD, EEF1E1, RPS6, GART, DNM2, RPS9, RPL13A, PRKAR1A, NME2, RPL24, LARS1, RPL34, AIMP1, PSAP, RPL14, RPS8, GAPDH, GSTP1, ATP5F1A, PAICS, RPS14, RPL23A, RPSA, TKT, RPS18, CD44, RPL6, GPI, SUMO2, RPL10, RPS24, SLC2A1, RPL32, EBP, VAPB, RPL37A, VAMP2, GMPS, RPL23, GC, RPS20, RPLP2, RPS25, RPS13, ENO2, SLC16A1, LDHA, RAN, RPL18, RPL28, RPL26, RPL17, RPL36, RPL27, RPS15, ACLY, RPS19, RPS28, RPS5, MARCKS, PSMC3, RPS10, PTGES3, PSMC6, PHGDH, RPS7, ENO1, RPL35A, ALDOA, RPL11, LPL, CTPS1, MTHFD1, RPS27, AK2 |
| Metabolism of amino acids and derivatives | HSA-71291 | 89 | 2,55E-45 | PSMC4, AHCY, RPL18A, RPL19, RPS12, RPL35, PSMA6, SLC7A5, RPL8, DARS1, PSMB6, RPS11, PSMA5, RPS27A, GLUD1, RPS3, RPL29, RPS23, CKB, RPL13, RPL15, RPL38, PSMD2, PSMC5, RPL4, RPS15A, RPL7, RPS2, SERINC1, RPL3, RPL36AL, RPL27A, RPL21, RPS3A, RPL22, ALDH9A1, RPS26, PSMB5, PSMD4, RPL5, RPL7A, RPS4X, RPL10A, SLC44A1, IARS1, SLC3A2, RPS16, EEF1E1, RPS6, RPS9, RPL13A, RPL24, LARS1, RPL34, AIMP1, RPL14, RPS8, RPS14, RPL23A, RPSA, RPS18, RPL6, RPL10, RPS24, RPL32, RPL37A, RPL23, RPS20, RPLP2, RPS25, RPS13, RPL18, RPL28, RPL26, RPL17, RPL36, RPL27, RPS15, RPS19, RPS28, RPS5, PSMC3, RPS10, PSMC6, PHGDH, RPS7, RPL35A, RPL11, RPS27 |
| Signal Transduction | HSA-162582 | 138 | 7,45E-10 | FKBP4, PSMC4, MYH9, CSNK2A1, PFN1, CPD, DDX5, HSPE1, CLTA, H4C6, TUBA4A, HSPB1, YWHAH, SNAP23, RAP1B, LMAN1, APOE, H3-3B, CCT7, TUBB2B, THBS1, ADAM10, LMNB1, PSMA6, GNAO1, YWHAE, ATP6V0A1, RAB7A, ESYT1, PSMB6, PSMA5, RPS27A, CCT6A, ACTC1, ATP6V0D1, FABP5, CKB, CCT2, SPINT2, H2AC8, FASN, CTNNA1, GNB2, YWHAG, KIF5B, H2AZ2, PSMD2, PSMC5, RAB6A, TLN1, RUVBL1, TUBB3, PPP2R1A, HNRNPM, SRRM1, P4HB, H2AC20, HSP90AA1, SFN, TUBB4B, HNRNPA1, FYN, FN1, RAC1, DYNC1H1, HNRNPH1, PTBP1, SFPQ, VCP, MYH10, COL6A1, PSMB5, PARP1, KIF14, F11R, S100A9, PSMD4, H2BC21, FLNA, GNAI3, MYO6, RBMX, GNAS, HSP90AB1, YBX1, YWHAB, RBBP4, RCC2, CSNK2B, GNB1, TUBB2A, RPS6, YWHAQ, DNM2, TFRC, CIT, PRKAR1A, DBN1, JUP, ACTN1, PSAP, YWHAZ, MYH11, CTNND1, FKBP1A, MDK, RHOA, CAPZB, RACGAP1, STMN1, DDX39B, TRA2B, VAPB, HDAC2, CFL1, CKAP5, TUBA1A, SLC1A5, VIM, MYL6, HNRNPC, MYL12B, PTPRS, H2BC14, PHB, CLTC, NCAM1, PSMC3, PTGES3, PSMC6, SPTAN1, MYH14, TPM4, ACTB, CTNNB1, CDC42, TPM3, RPS27 |
| Cellular responses to stimuli | HSA-8953897 | 136 | 9,68E-57 | FKBP4, PSMC4, ST13, CSNK2A1, RPL18A, RPL19, RPS12, CAT, H1-3, H4C6, TUBA4A, RPA1, H3-3B, EIF2S1, RPL35, TUBB2B, LMNB1, PSMA6, RPL8, PRDX1, CAPZA1, YWHAE, PRDX5, PSMB6, RPS11, PSMA5, RPS27A, RPS3, ATP6V0D1, RPL29, ALB, RPS23, HSPA9, HSP90B1, GCN1, PRDX2, H2AC8, H1-4, RPL13, H2AZ2, RPL15, RPL38, PSMD2, PSMC5, RPL4, TLN1, RPS15A, TUBB3, CALR, TALDO1, HSPA5, P4HB, H1-5, H2AC20, HSP90AA1, EEF1A1, H1-2, RPL7, TUBB4B, RPS2, H1-0, RPL3, RPL36AL, RPL27A, RPL21, RPS3A, RPL22, RPS26, DYNC1H1, VCP, PSMB5, CSRP1, LMNA, PSMD4, H2BC21, RPL5, HSP90AB1, RPL7A, RBBP4, RPS4X, RPL10A, HSPA1B, CSNK2B, RPS16, TUBB2A, RPS6, RPS9, RPL13A, RPL24, RPL34, HSPA2, RPL14, RPS8, KHSRP, GSTP1, PDIA6, RPS14, RPL23A, RPSA, TKT, RPS18, HMGA1, CAPZB, RPL6, RPL10, RPS24, RPL32, RPL37A, RPL23, RPS20, RPLP2, RPS25, RPS13, HSPA8, TUBA1A, RPL18, RPL28, RPL26, RPL17, RPL36, RPL27, RPS15, RPS19, RPS28, RPS5, H2BC14, PSMC3, RPS10, PTGES3, PSMC6, HBB, RPS7, RPL35A, RPL11, MAP1LC3B, RPS27 |
| Cellular responses to stress | HSA-2262752 | 135 | 9,68E-57 | FKBP4, PSMC4, ST13, CSNK2A1, RPL18A, RPL19, RPS12, CAT, H1-3, H4C6, TUBA4A, RPA1, H3-3B, EIF2S1, RPL35, TUBB2B, LMNB1, PSMA6, RPL8, PRDX1, CAPZA1, YWHAE, PRDX5, PSMB6, RPS11, PSMA5, RPS27A, RPS3, ATP6V0D1, RPL29, ALB, RPS23, HSPA9, HSP90B1, GCN1, PRDX2, H2AC8, H1-4, RPL13, H2AZ2, RPL15, RPL38, PSMD2, PSMC5, RPL4, TLN1, RPS15A, TUBB3, CALR, TALDO1, HSPA5, P4HB, H1-5, H2AC20, HSP90AA1, EEF1A1, H1-2, RPL7, TUBB4B, RPS2, H1-0, RPL3, RPL36AL, RPL27A, RPL21, RPS3A, RPL22, RPS26, DYNC1H1, VCP, PSMB5, LMNA, PSMD4, H2BC21, RPL5, HSP90AB1, RPL7A, RBBP4, RPS4X, RPL10A, HSPA1B, CSNK2B, RPS16, TUBB2A, RPS6, RPS9, RPL13A, RPL24, RPL34, HSPA2, RPL14, RPS8, KHSRP, GSTP1, PDIA6, RPS14, RPL23A, RPSA, TKT, RPS18, HMGA1, CAPZB, RPL6, RPL10, RPS24, RPL32, RPL37A, RPL23, RPS20, RPLP2, RPS25, RPS13, HSPA8, TUBA1A, RPL18, RPL28, RPL26, RPL17, RPL36, RPL27, RPS15, RPS19, RPS28, RPS5, H2BC14, PSMC3, RPS10, PTGES3, PSMC6, HBB, RPS7, RPL35A, RPL11, MAP1LC3B, RPS27 |
| Signaling by ROBO receptors | HSA-376176 | 83 | 3,60E-55 | PSMC4, RPL18A, RPL19, PFN1, RPS12, VASP, RPL35, PSMA6, RPL8, GPC1, PSMB6, RPS11, PSMA5, RPS27A, RPS3, RPL29, RPS23, RPL13, RPL15, RPL38, PSMD2, PSMC5, RPL4, PABPC1, RPS15A, RPL7, RPS2, RPL3, RPL36AL, RPL27A, RPL21, RPS3A, RPL22, RAC1, RPS26, PSMB5, PSMD4, RPL5, RPL7A, RPS4X, RPL10A, RPS16, RPS6, RPS9, RPL13A, RPL24, RPL34, RPL14, RPS8, RPS14, RPL23A, RPSA, RPS18, RHOA, RPL6, RPL10, RPS24, RPL32, RPL37A, RPL23, RPS20, RPLP2, RPS25, RPS13, RPL18, RPL28, RPL26, RPL17, RPL36, RPL27, RPS15, RPS19, RPS28, RPS5, PSMC3, RPS10, PSMC6, RPS7, RPL35A, RPL11, EIF4A3, CDC42, RPS27 |
| Signaling by Rho GTPases | HSA-194315 | 78 | 9,63E-21 | MYH9, PFN1, CPD, HSPE1, H4C6, TUBA4A, YWHAH, SNAP23, LMAN1, H3-3B, CCT7, TUBB2B, LMNB1, YWHAE, RAB7A, ESYT1, CCT6A, ACTC1, CKB, CCT2, H2AC8, CTNNA1, YWHAG, KIF5B, H2AZ2, TUBB3, PPP2R1A, SRRM1, H2AC20, HSP90AA1, SFN, TUBB4B, RAC1, DYNC1H1, VCP, MYH10, KIF14, S100A9, H2BC21, FLNA, MYO6, RBMX, HSP90AB1, YWHAB, RCC2, TUBB2A, YWHAQ, TFRC, CIT, DBN1, JUP, ACTN1, YWHAZ, MYH11, RHOA, CAPZB, RACGAP1, DDX39B, TRA2B, VAPB, CFL1, CKAP5, TUBA1A, SLC1A5, VIM, MYL6, HNRNPC, MYL12B, H2BC14, CLTC, SPTAN1, MYH14, TPM4, ACTB, CTNNB1, CDC42, TPM3, RPS27 |
| Immune System | HSA-168256 | 137 | 1,67E-17 | PSMC4, MIF, MYH9, CHGA, VTN, LTF, CAT, CLTA, VASP, CANX, TUBA4A, SNAP23, RAP1B, KRT1, SNRPA1, SNAP25, TUBB2B, KIF23, ADAM10, LYZ, LMNB1, UBE2K, PSMA6, CRISPLD2, ERP44, CAPZA1, CD81, ATP6V0A1, RAB10, RAB7A, PSMB6, PSMA5, RPS27A, FTH1, AHSG, CXADR, CCT8, KPNB1, ATP6V0D1, EIF4A1, HSPA9, FABP5, INPPL1, CCT2, HSP90B1, PDIA3, PA2G4, KIF5B, EEF2, PSMD2, PSMC5, RAB6A, PRKDC, TOLLIP, PKM, TUBB3, CALR, TALDO1, HSPA5, PPP2R1A, P4HB, LAMP1, HSP90AA1, UBA1, TUBB, EEF1A1, TUBB4B, HMGB1, VAPA, ANXA2, FYN, HNRNPA2B1, FN1, RAC1, DYNC1H1, IGF2R, VCP, XRCC6, ILF2, PSMB5, DHX9, S100A9, HRNR, PSMD4, FLNA, PGAM1, HSP90AB1, TSPAN14, YWHAB, RAB14, KIF4A, HSPA1B, CSNK2B, APRT, TXNDC5, GDI2, TUBB2A, FSCN1, DNM2, XRCC5, NME2, JUP, PSAP, YWHAZ, GSTP1, FKBP1A, CD44, HLA-B, RHOA, CAPZB, SIGIRR, RACGAP1, GPI, DCD, VAMP2, PPIA, FLNB, CFL1, HSPA8, TUBA1A, VIM, RAB5C, PRKCSH, ACLY, CLTC, NCAM1, PSMC3, PSMC6, SPTAN1, DDX3X, HBB, ACTB, CTNNB1, ALDOA, EIF4A3, CDC42, CD59 |
| Infectious disease | HSA-5663205 | 166 | 1,64E-70 | FKBP4, PSMC4, SNRPD3, MYH9, SUPT16H, RPL18A, RPL19, DDX5, RPS12, LTF, CLTA, H4C6, CANX, TUBA4A, YWHAH, TRIM28, SNAP25, RPL35, TUBB2B, PPIG, PSMA6, RPL8, GPC1, YWHAE, RAB7A, SNRPF, PSMB6, RPS11, PSMA5, RPS27A, SSRP1, RPS3, KPNB1, RPL29, RPS23, NPM1, PPIB, H2AC8, GNB2, YWHAG, RPL13, EEF2, RPL15, RPL38, PSMD2, PSMC5, RPL4, RPS15A, TUBB3, CALR, H2AC20, H2AC21, HSP90AA1, TUBB, EEF1A1, RPL7, GANAB, SFN, TUBB4B, HNRNPA1, RPS2, SNRPD2, RPL3, RPL36AL, RPL27A, RPL21, RPS3A, RPL22, FYN, RAC1, RPS26, DYNC1H1, SFPQ, VCP, XRCC6, PSMB5, PARP1, VTA1, ATP1B1, PSMD4, H2BC21, GNAI3, RPL5, GNAS, HSP90AB1, SLC25A5, RPL7A, SDC4, YWHAB, PGK1, RBBP4, RPS4X, RPL10A, HSPA1B, HNRNPK, RPS16, GNB1, TUBB2A, RPS6, PSIP1, YWHAQ, CD9, RPS9, RPL13A, XRCC5, PRKAR1A, RPL24, RPL34, YWHAZ, RPL14, RPS8, CTNND1, FKBP1A, RPS14, RPL23A, RPSA, RPS18, SNRPG, HLA-B, HMGA1, SNRPE, RPL6, SNRPB, RPL10, RPS24, RPL32, RPL37A, VAMP2, PPIA, RPL23, RPS20, HDAC2, RPLP2, BANF1, RPS25, RPS13, TUBA1A, ATP1A3, ATP1A1, RAN, RPL18, RPL28, RPL26, RPL17, RPL36, RPL27, RPS15, PRKCSH, RPS19, RPS28, RPS5, H2BC14, CLTC, PSMC3, RPS10, PTGES3, PSMC6, ACTB, RPS7, CTNNB1, ENO1, RPL35A, RPL11, CDC42, MAP1LC3B, RPS27 |
| Innate Immune System | HSA-168249 | 99 | 9,51E-21 | PSMC4, MIF, MYH9, CHGA, VTN, LTF, CAT, SNAP23, RAP1B, KRT1, SNAP25, ADAM10, LYZ, UBE2K, PSMA6, CRISPLD2, ERP44, CAPZA1, CD81, ATP6V0A1, RAB10, RAB7A, PSMB6, PSMA5, RPS27A, FTH1, AHSG, CCT8, KPNB1, ATP6V0D1, FABP5, CCT2, HSP90B1, PA2G4, EEF2, PSMD2, PSMC5, RAB6A, PRKDC, TOLLIP, PKM, PPP2R1A, LAMP1, HSP90AA1, TUBB, EEF1A1, TUBB4B, HMGB1, VAPA, ANXA2, FYN, RAC1, DYNC1H1, IGF2R, VCP, XRCC6, ILF2, PSMB5, DHX9, S100A9, HRNR, PSMD4, PGAM1, HSP90AB1, TSPAN14, RAB14, HSPA1B, CSNK2B, APRT, TXNDC5, GDI2, DNM2, XRCC5, NME2, JUP, PSAP, GSTP1, CD44, HLA-B, RHOA, SIGIRR, GPI, DCD, PPIA, CFL1, HSPA8, RAB5C, PRKCSH, ACLY, PSMC3, PSMC6, SPTAN1, DDX3X, HBB, ACTB, CTNNB1, ALDOA, CDC42, CD59 |
| Disease | HSA-1643685 | 195 | 6,61E-56 | FKBP4, PSMC4, SNRPD3, MYH9, SUPT16H, AHCY, RPL18A, RPL19, DDX5, RPS12, LTF, MSH6, CLTA, H4C6, CANX, TUBA4A, YWHAH, RAP1B, TRIM28, RPA1, H3-3B, SNAP25, RPL35, TUBB2B, THBS1, ADAM10, PPIG, LMNB1, PSMA6, RPL8, PRDX1, GPC1, YWHAE, RAB7A, SNRPF, PSMB6, RPS11, PSMA5, RPS27A, SSRP1, RPS3, KPNB1, RPL29, RPS23, NPM1, PPIB, PRDX2, H2AC8, GNB2, YWHAG, KIF5B, RPL13, EEF2, H2AZ2, RPL15, RPL38, PSMD2, PSMC5, RPL4, TLN1, RPS15A, TUBB3, CALR, TALDO1, PPP2R1A, EEF1G, H2AC20, H2AC21, BSG, HSP90AA1, TUBB, EEF1A1, RPL7, GANAB, SFN, TUBB4B, HNRNPA1, RPS2, SNRPD2, HMGB1, RPL3, RPL36AL, RPL27A, RPL21, RPS3A, RPL22, FYN, SND1, FN1, RAC1, RPS26, DYNC1H1, SFPQ, VCP, XRCC6, PSMB5, PARP1, VTA1, ATP1B1, LMNA, S100A9, PSMD4, H2BC21, GNAI3, RPL5, GNAS, HSP90AB1, SLC25A5, RPL7A, SDC4, YWHAB, PGK1, RBBP4, RPS4X, RPL10A, HSPA1B, HNRNPK, SLC3A2, APRT, RPS16, GNB1, TUBB2A, RPS6, PSIP1, YWHAQ, CD9, RPS9, RPL13A, XRCC5, PRKAR1A, RPL24, RPL34, YWHAZ, RPL14, RPS8, CTNND1, FKBP1A, RPS14, RPL23A, RPSA, RPS18, SNRPG, HLA-B, HMGA1, SNRPE, RPL6, SNRPB, RPL10, RPS24, SLC2A1, RPL32, RPL37A, VAMP2, PPIA, RPL23, RPS20, HDAC2, RPLP2, BANF1, RPS25, RPS13, TUBA1A, SLC16A1, ATP1A3, ATP1A1, RAN, RPL18, RPL28, RPL26, RPL17, RPL36, RPL27, RPS15, PRKCSH, RPS19, RPS28, RPS5, H2BC14, PHB, CLTC, PSMC3, RPS10, PTGES3, PSMC6, TPM4, ACTB, RPS7, CTNNB1, ENO1, RPL35A, RPL11, CDC42, MAP1LC3B, TPM3, RPS27 |
| Neutrophil degranulation | HSA-6798695 | 71 | 2,03E-24 | MIF, LTF, CAT, SNAP23, RAP1B, KRT1, SNAP25, ADAM10, LYZ, CRISPLD2, ERP44, ATP6V0A1, RAB10, RAB7A, PSMA5, FTH1, AHSG, CCT8, KPNB1, FABP5, CCT2, PA2G4, EEF2, PSMD2, RAB6A, TOLLIP, PKM, LAMP1, HSP90AA1, TUBB, EEF1A1, TUBB4B, HMGB1, VAPA, ANXA2, RAC1, DYNC1H1, IGF2R, VCP, XRCC6, ILF2, S100A9, HRNR, PGAM1, HSP90AB1, TSPAN14, RAB14, HSPA1B, CSNK2B, APRT, TXNDC5, GDI2, XRCC5, NME2, JUP, PSAP, GSTP1, CD44, HLA-B, RHOA, GPI, PPIA, HSPA8, RAB5C, ACLY, PSMC3, SPTAN1, DDX3X, HBB, ALDOA, CD59 |
| Nervous system development | HSA-9675108 | 119 | 3,60E-55 | PSMC4, MYH9, CSNK2A1, RPL18A, RPL19, PFN1, RPS12, CLTA, VASP, TUBA4A, RPL35, TUBB2B, ADAM10, PSMA6, RPL8, GPC1, PSMB6, RPS11, PSMA5, RPS27A, RPS3, RPL29, RPS23, RPL13, RPL15, RPL38, PSMD2, PSMC5, RPL4, PABPC1, TLN1, RPS15A, TUBB3, CRMP1, HSP90AA1, EZR, RPL7, TUBB4B, RPS2, DPYSL3, RPL3, RPL36AL, RPL27A, RPL21, RPS3A, RPL22, FYN, RAC1, RPS26, MYH10, COL6A1, PSMB5, PSMD4, RPL5, HSP90AB1, RPL7A, RPS4X, RPL10A, KIF4A, CSNK2B, RPS16, TUBB2A, RPS6, DNM2, EPHA6, RPS9, RPL13A, RPL24, RPL34, RPL14, MYH11, RPS8, MBP, RPS14, RPL23A, RPSA, RPS18, RHOA, RPL6, RPL10, RPS24, RPL32, RPL37A, RPL23, DPYSL2, RPS20, HDAC2, RPLP2, CFL1, RPS25, RPS13, HSPA8, TUBA1A, MYL6, RPL18, RPL28, MYL12B, RPL26, RPL17, RPL36, RPL27, RPS15, RPS19, RPS28, RPS5, CLTC, NCAM1, PSMC3, RPS10, PSMC6, SPTAN1, MYH14, ACTB, RPS7, RPL35A, RPL11, EIF4A3, CDC42, RPS27 |
| Axon guidance | HSA-422475 | 117 | 3,54E-55 | PSMC4, MYH9, CSNK2A1, RPL18A, RPL19, PFN1, RPS12, CLTA, VASP, TUBA4A, RPL35, TUBB2B, ADAM10, PSMA6, RPL8, GPC1, PSMB6, RPS11, PSMA5, RPS27A, RPS3, RPL29, RPS23, RPL13, RPL15, RPL38, PSMD2, PSMC5, RPL4, PABPC1, TLN1, RPS15A, TUBB3, CRMP1, HSP90AA1, EZR, RPL7, TUBB4B, RPS2, DPYSL3, RPL3, RPL36AL, RPL27A, RPL21, RPS3A, RPL22, FYN, RAC1, RPS26, MYH10, COL6A1, PSMB5, PSMD4, RPL5, HSP90AB1, RPL7A, RPS4X, RPL10A, KIF4A, CSNK2B, RPS16, TUBB2A, RPS6, DNM2, EPHA6, RPS9, RPL13A, RPL24, RPL34, RPL14, MYH11, RPS8, RPS14, RPL23A, RPSA, RPS18, RHOA, RPL6, RPL10, RPS24, RPL32, RPL37A, RPL23, DPYSL2, RPS20, RPLP2, CFL1, RPS25, RPS13, HSPA8, TUBA1A, MYL6, RPL18, RPL28, MYL12B, RPL26, RPL17, RPL36, RPL27, RPS15, RPS19, RPS28, RPS5, CLTC, NCAM1, PSMC3, RPS10, PSMC6, SPTAN1, MYH14, ACTB, RPS7, RPL35A, RPL11, EIF4A3, CDC42, RPS27 |
| Developmental Biology | HSA-1266738 | 152 | 5,86E-51 | PSMC4, KRT14, MYH9, CSNK2A1, RPL18A, RPL19, PFN1, RPS12, CLTA, H4C6, VASP, KRT9, TUBA4A, KRT31, KRT33B, KRT5, KRT1, KRT6B, H3-3B, KRT85, RPL35, TUBB2B, ADAM10, PSMA6, RPL8, GPC1, KRT10, PSMB6, RPS11, PSMA5, RPS27A, RPS3, RPL29, RPS23, KRT16, H2AC8, CTNNA1, RPL13, H2AZ2, KRT17, RPL15, RPL38, PSMD2, PSMC5, KRT2, RPL4, PABPC1, TLN1, RPS15A, TUBB3, CRMP1, KRT7, H2AC20, HSP90AA1, EZR, RPL7, TUBB4B, RPS2, DPYSL3, RPL3, RPL36AL, RPL27A, RPL21, RPS3A, RPL22, THRAP3, FYN, RAC1, RPS26, MYH10, KRT19, COL6A1, PSMB5, PSMD4, H2BC21, RPL5, HSP90AB1, RPL7A, RBBP4, RPS4X, RPL10A, KIF4A, CSNK2B, RPS16, KRT6A, TUBB2A, RPS6, KRT18, DNM2, EPHA6, RPS9, RPL13A, JUP, KRT34, RPL24, RPL34, RPL14, MYH11, RPS8, MBP, RPS14, RPL23A, RPSA, RPS18, RHOA, RPL6, RPL10, RPS24, RPL32, RPL37A, RPL23, DPYSL2, RPS20, HDAC2, RPLP2, CFL1, RPS25, RPS13, HSPA8, TUBA1A, KRT86, MYL6, RPL18, KRT8, RPL28, MYL12B, RPL26, RPL17, RPL36, RPL27, RPS15, RPS19, RPS28, RPS5, H2BC14, CLTC, NCAM1, PSMC3, RPS10, PSMC6, SPTAN1, MYH14, ACTB, RPS7, CTNNB1, RPL35A, RPL11, EIF4A3, LPL, CDC42, DEK, RPS27 |
