## Supplementary Table 3 for "Multi-omics characterization of extracellular vesicles derived from virus-positive Merkel cell carcinoma cells"

**Supplementary Table S3.** Twenty-five most abundant miRNAs identified in WaGa and MKL-1 cells and their corresponding extracellular vesicles.

| **Rank** | **WaGa cells** | **WaGa EVs** | **MKL-1 cells** | **MKL-1 EVs** |
| --- | --- | --- | --- | --- |
| 1 | hsa-let-7a-5p | hsa-miR-10b-5p | hsa-let-7b-5p | hsa-let-7a-5p |
| 2 | hsa-miR-10b-5p | hsa-miR-375 | hsa-miR-375 | hsa-miR-375 |
| 3 | hsa-miR-375 | hsa-let-7b-5p | hsa-miR-486-5p | hsa-miR-10b-5p |
| 4 | hsa-let-7b-5p | hsa-let-7a-5p | hsa-miR-92a-3p | hsa-let-7b-5p |
| 5 | hsa-let-7f-5p | hsa-miR-182-5p | hsa-miR-200c-3p | hsa-miR-486-5p |
| 6 | hsa-miR-182-5p | hsa-miR-320a | hsa-miR-181a-5p | hsa-miR-182-5p |
| 7 | hsa-miR-30a-5p | hsa-miR-378a-3p | hsa-miR-93-5p | hsa-let-7f-5p |
| 8 | hsa-miR-26a-5p | hsa-miR-191-5p | hsa-miR-103a-3p | hsa-miR-320a |
| 9 | hsa-let-7g-5p | hsa-miR-423-5p | hsa-miR-320a | hsa-miR-30a-5p |
| 10 | hsa-miR-30d-5p | hsa-miR-30a-5p | hsa-miR-30d-5p | hsa-miR-30d-5p |
| 11 | hsa-let-7i-5p | hsa-miR-92a-3p | hsa-miR-92b-3p | hsa-miR-92a-3p |
| 12 | hsa-miR-92a-3p | hsa-let-7f-5p | hsa-miR-182-5p | hsa-miR-25-3p |
| 13 | hsa-miR-103a-3p | hsa-miR-30d-5p | hsa-let-7g-5p | hsa-miR-191-5p |
| 14 | hsa-miR-93-5p | hsa-miR-486-5p | hsa-let-7i-5p | hsa-miR-92b-3p |
| 15 | hsa-miR-148a-3p | hsa-miR-26a-5p | hsa-miR-10b-5p | hsa-miR-98-5p |
| 16 | hsa-miR-200c-3p | hsa-miR-25-3p | hsa-miR-378a-3p | hsa-miR-423-5p |
| 17 | hsa-miR-486-5p | hsa-miR-93-5p | hsa-miR-191-5p | hsa-miR-181a-5p |
| 18 | hsa-miR-16-5p | hsa-miR-181a-5p | hsa-let-7f-5p | hsa-miR-378a-3p |
| 19 | hsa-miR-191-5p | hsa-miR-885-3p | hsa-miR-30a-5p | hsa-let-7i-5p |
| 20 | hsa-miR-1-3p | hsa-miR-320b | hsa-miR-181b-5p | hsa-miR-26a-5p |
| 21 | hsa-miR-9-5p | hsa-miR-148a-3p | hsa-miR-744-5p | hsa-miR-30c-5p |
| 22 | hsa-miR-181a-5p | hsa-let-7i-5p | hsa-miR-423-3p | hsa-let-7g-5p |
| 23 | hsa-miR-25-3p | hsa-miR-146b-5p | hsa-miR-26a-5p | hsa-miR-1246 |
| 24 | hsa-miR-26b-5p | hsa-miR-30c-5p | hsa-miR-423-5p | hsa-miR-103a-3p |
| 25 | hsa-miR-98-5p | hsa-miR-423-3p | hsa-miR-196b-5p | hsa-miR-744-5p |
